# Developing a Coarse-Grained Model for Bacterial Cell Walls and Evaluating Mechanical Properties and Free Energy Barriers

**DOI:** 10.1101/2020.03.12.985218

**Authors:** Rakesh Vaiwala, Pradyumn Sharma, Mrinalini Puranik, K. Ganapathy Ayappa

## Abstract

The bacterial cell envelope of Gram-negative bacteria is a complex biological barrier with multiple layers consisting of the inner membrane, periplasm of peptidoglycan and the outer membrane with lipopolysaccharides (LPS). With rising antimicrobial resistance there is increasing interest in understanding interactions of small molecules with the cell membrane to aid in the development of novel drug molecules. Hence suitable representations of the bacterial membrane are required to carry out meaningful molecular dynamics simulations. Given the complexity of the cell envelope, fully atomistic descriptions of the cell membrane with explicit solvent are computationally prohibitive, allowing limited sampling with small system sizes. However coarse-grained (CG) models such as MARTINI allow one to study phenomena at physiologically relevant length and time scales. Although MARTINI models for lipids and the LPS are available in literature, a suitable CG model of peptidoglycan is lacking. In this manuscript we develop a CG model of the peptidoglycan network within the MARTINI framework using an all-atom model developed by Gumbart et al. ^1^. The model is parametrized to reproduce the structural properties of the glycan strands, such as the end-to-end distance, equilibrium angle between adjacent peptides along the strands and area per disaccharide. Mechanical properties such as the area compressibility and the bending modulus are accurately reproduced. While developing novel antibiotics it is important to assess barrier properties of the peptidogylcan network. We evaluate and compare the free energy of insertion for a thymol molecule using umbrella sampling on both the MARTINI and all-atom peptidoglycan models. The insertion free energy was found to be less than k_B_T for both the MARTINI and all-atom models. Additional restraint free simulations reveal rapid translocation of thymol across peptidogylcan. We expect that the proposed MARTINI model for peptidoglycan will be useful in understanding phenomena associated with bacterial cell walls at larger length and time scales, thereby overcoming the current limitations of all-atom models.

## 1 Introduction

Bacterial cells are surrounded by cell walls with a distinct architecture. In Gram-negative bacteria, cells are protected by an inner phospholipid bilayer membrane, a periplasmic space containing a cell wall made up of peptidoglycan, and an outer membrane consisting of asymmetric leaflets of phospholipids and lipopolysaccharides.^2^ In the absence of the outer membrane, the cell wall for Gram-positive bacteria, consists of a relatively thick peptidogylcan structure. Peptidoglycan, also known as murein, is the main stress-bearing component of bacterial cells, resisting internal turgor pressure and determining the eventual shape of the cell. A molecular understanding of the bacterial cell membrane and its components play a key role in development of persistence in antimicrobial resistance (AMR) bacterial strains. Several antibiotics directly target various components of the cell membrane; for example colistin targets lipopolysaccharides,^3^ vancomycin inhibits synthesis pathways of peptidoglycan,^4^ and lysins enzymatically degrade peptidoglycan.^5^ In this manuscript our primary focus lies in developing a coarse grained model for peptidoglycan.

In its architecture, peptidoglycan is a mesh-like structure constructed by oligomeric strands of glycans, which are cross-linked by stems of peptides (muropeptides). The glycan strands are polymers made up of alternating units of N-acetylglucosamine (NAG) and N-acetylmuramic acid (NAM). A short peptide containing 5 amino acids is attached to lactyl moiety of each NAM residue. ^6^ The usual sequence of the amino acids in the penta-peptide is: L-alanine (L-ALA), D-iso-glutamate (GLU), meso-diamino pimelic acid (DAP), D-alanine (D-ALA) and D-alanine (D-ALA).^7^ The molecular structures of these sugar units and the peptide forming amino acids are illustrated in Figure 1(A). In general, the periplasm containing peptidoglycan is multilayered,^8^ with the exception of E. coli which has more than 75% of glycan strands arranged in a monolayer^9^ with a thickness of 4 nm.^10^ Glycan strands show a broad distribution of lengths, with a mean length ranging from 21 to 40 disaccharides of NAG and NAM residues.^11,12^ A single disaccharide (1-mer unit) is of length 1.03 nm. ^13^

**Figure 1:**
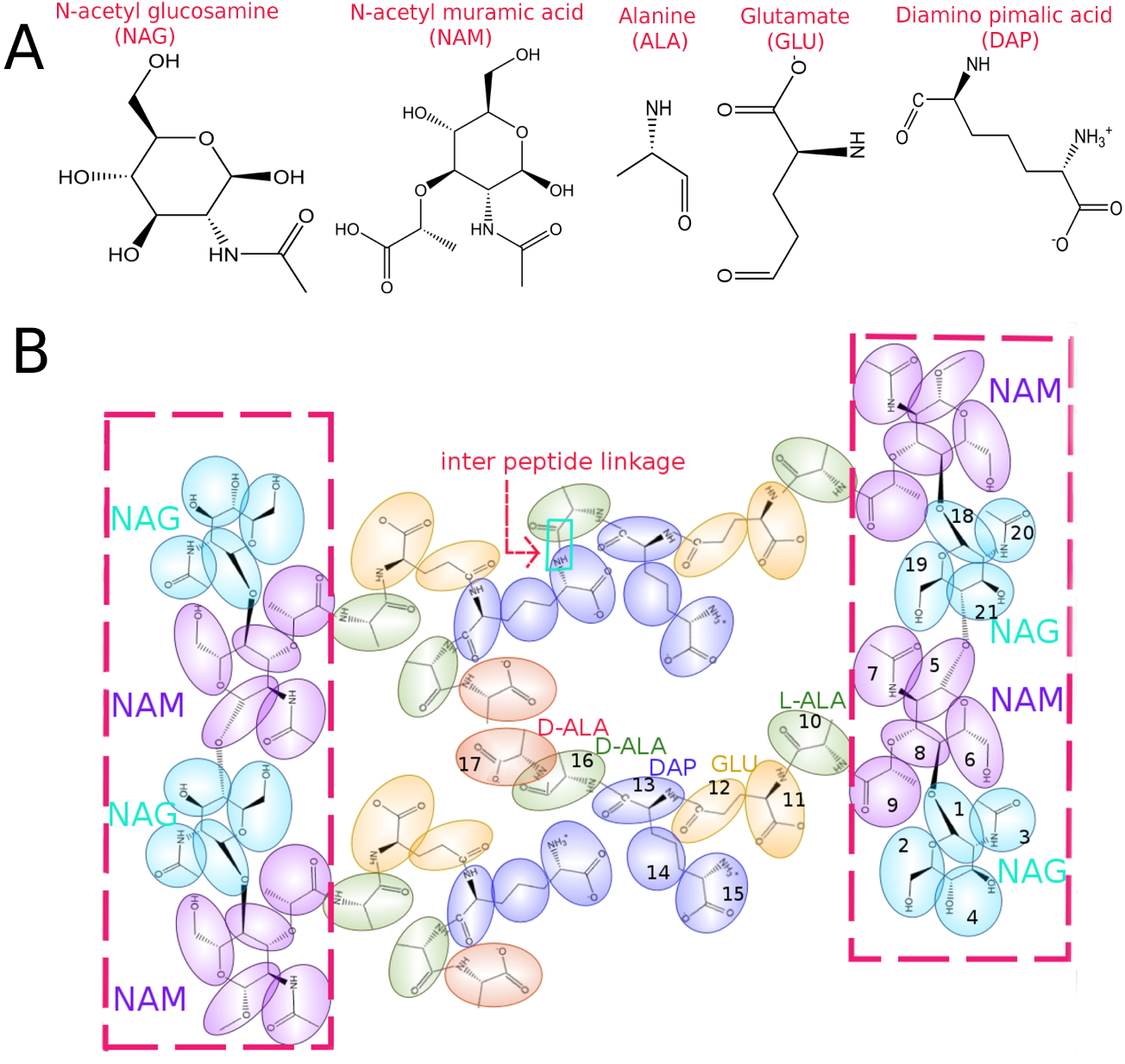
(A) Atomic structures of the molecular building blocks in construction of peptidoglycan. (B) Schematic showing an atomic structure of peptidoglycan strands (each 2-mer long) and coarse-grained mapping scheme with tinted MARTINI beads superimposed upon the underlying atomic structure. Coarse graining from the all-atom structure to the MAR-TINI model results in about 7 fold reduction in the number of atoms. Each mer is represented by 17 beads in the MARTINI force field, and the numeric indicates the bead labels. The glycan strands are shown within the boxes, and the ordering of amino acids from a sugar backbone is L-ALA, GLU, DAP, D-ALA and D-ALA. As indicated by a small rectangle, a pair of peptides is covalently bonded via inter peptide linkage to form a cross-link between the glycan strands. While linking the peptides, the terminal residue D-ALA in one of the peptides is eliminated.

To understand the organization of the glycan strands relative to the cell surface, two diverse models were proposed based on electron microscopy experiments.^14,15^ According to the ‘circumferential layered’ model, ^14^ glycan strands orient parallel to the cell surface. In contrast, the ‘scaffold’ model assumes that the strands protrude out from the cytoplasmic membrane perpendicular to the cell surface.^15^ The relatively thin (≤ 4 nm) E. coli cell surface and the orientation of wrinkles in electron tomography data^10^ support the circumferential model which has been widely used while developing models for peptidogylcan.^1,16,17^ Several models for peptidoglycan ranging from all-atom to supra coarse-grained have been developed in the literature, and we review them next.

Using energy minimization in an all-atom model Leps et al. ^18^ computed energetically favourable conformations of the glycan backbone and found that the backbone adopts extended structures with each disaccharide spanning 0.98-1.02 nm in length. The lactyl sites in NAM orient away from the murein backbone, and the propagation angle between subsequent sites is found to be 80-100°. Koch ^19^ extended the conformational analysis for a penta-muropeptide, and a nona-muropeptide in aqueous and cytoplasmic environments. A three dimensional structure of bacterial cell wall was proposed by Meroueh et al. ^20^ using molecular dynamics simulations on a network formed by 8-mer long peptidoglycan strands. The peptides adopted three-fold symmetry of orientation along the glycan helices, with a minimum pore diameter of 7 nm in the mesh structure. In an atomistic MD simulation of peptidoglycan Gumbart et al. ^1^ capture the monolayer thickness, pore size, and anisotropic elasticity in two orthogonal directions of glycans and peptides in general agreement with experimental data, supporting the disordered circumferential model of peptidoglycan. The model exhibits strain stiffening behavior at moderate to high surface tension. ^21^ In other studies, stable interactions of lysozyme with glycan reveal that O-acetylated glycan is highly distorted, disrupting interactions with lysozyme. ^22^ An atomistic model of peptidoglycan compatible with a CHARMM36 force field was employed recently by Kim et al. ^23^ to investigate structural rearrangement of nascent peptidoglycan in presence of penicillin-binding protein (PBP1b).

In addition to atomistic and coarse grained molecular descriptions for bacterial cell walls, a few models which permit studying deformations of the entire cell wall have been developed. Huang et al. ^16^ developed a model for peptidoglycan using a network of springs to capture the mechanical response of Escherichia coli cells. Incorporating forces due to springs, bending, and the osmotic pressure difference at vertices formed by peptides and glycan springs. The mechanical response of cell shape to cell wall damage was predicted. A supra coarse-grained model of the cell wall sacculus was developed by Nguyen et al. ^17^ to study remodeling of peptidoglycan during bio-synthesis and growth of the cell wall. With local coordination of enzymes, the model sacculus prevents local defects caused by new material introduced via transpeptidation and transglycosylation, enabling enzymes to move along the glycan hoops, and thereby maintaining the cell wall integrity and rod-like shape.

Syma Khalid and co-workers^24–26^ have developed several models for the outer membrane of Gram-negative bacteria using united atom as well as MARTINI representations. Using united atom simulations Samsudin et al. ^27^ reveal the manner in which the distance between outer membrane porins such as OmpA and peptidoglycan is reduced due to binding of OmpA C-terminus residues and the Braun’s lipoprotein with peptidoglycan. Additionally binding of the C-terminus residues was found to assist the dimerization of OmpA in the absence of Braun’s lipoprotein. In a more complex model of the membrane by Boags et al. ^28^, which included peptidoglycan, the inner membrane with TolR protein, and the outer membrane with an OmpA dimer, the authors illustrate the role of non-covalent interactions in positioning the peptidoglycan layer in the presence of the TolR protein.

Simulating a model bacterial cell envelope with atomic details is computationally demanding, especially when time and length scales involved in membrane-associated collective phenomena are order of milliseconds and micrometers, respectively. Under such circumstances, coarse-grained (CG) models of the bacterial cell are ideally suited for large scale molecular simulations, allowing relatively larger systems to be investigated over longer time scales. Therefore, there exists a variety of coarse-grained models for lipids, amino acids, carbohydrates and nucleic acids. The methodology to devise the coarse-grained model parameters ranges from solvent-free models to more realistic explicit models that include chemical specificity.

MARTINI force fields for coarse-grained simulations were originally developed for lipids and cholesterol.^29^ Subsequently the MARTINI force field was extended to carbohydrates, ^30^ proteins^31^ and nucleotides.^32,33^ The bonded parameters for monosaccharides such as glucose and fructose, and disaccharides such as sucrose, maltose and cellobiose are optimized to match the conformations obtained from AA simulations. The CG models of 20 amino acids are systematically parametrized using 2000 proteins from Protein Data Bank.^31^ Following the MARTINI philosophy, López et al. ^34^ proposed CG models for more than 5 types of glycolipids. The model membranes of glycolipids are tested for their structural properties such as electron density, area per lipid and the membrane thickness. The MARTINI models are not without drawbacks.^35^ For instance, MARTINI sugars are sticky and form aggregates at concentrations below their solubility limits.^36^ The protein-protein interactions are unrealistic, giving rise to excessive free energies for protein dimerization. ^37^ By scaling down the well-depth of the Lennard-Jones interactions, the exaggerated clustering of proteins and aggregation of sugars can be avoided.^36,38^

Here we develop a MARTINI model for peptidogylcan by using an atomistic model we devised following Gumbart and co-workers.^1^ Coarse graining is carried out at various levels of increasing complexity in order to develop a robust MARTINI model for a peptidoglycan chain and a peptidoglycan network. At each level mapping is systematically carried out with the reference all-atom simulations in explicit water. Using a peptidoglycan network consisting of 21 glycan strands which are equivalent to 0.5 million atoms in all-atom simulations, we evaluate the stretch modulus, bending modulus and density distributions. Although there are recent studies attempting to understand antimicrobial activity of small molecules with the complex outer membrane of bacteria,^39–41^ the interaction of peptidoglycan with molecules has not been reported. To our knowledge, this is the first study to investigate the interactions of a small molecule with peptidoglycan. Potential of mean force computations carried out using MARTINI and all-atoms models illustrate the low barrier for translocation through the peptidoglycan layer.

The manuscript is organized as follows. The simulation methodology is detailed in Section 2. The mapping procedure and model validation are delineated in Section 3. This includes a discussion on potential of mean force between peptidoglycan and thymol molecule.

## 2 Simulation Methods

Following the bottom up approach, a MARTINI model of peptidoglycan is devised by mapping the distributions for bonds, angles and dihedrals on the target distributions, which are derived from the virtual CG-trajectories obtained in all-atom simulations. To develop the CG model in a systematic manner, we have carried out all-atom and MARTINI simulations on systems with a single glycan strand in water, a peptidoglycan chain containing peptides in water, and the peptidoglycan networks as depicted in Figure 2. All simulations are performed on GROMACS package,^42^ version 5.1.5, at 310 K and 1 bar pressure unless stated otherwise. The system size, number of solvent molecules and glycan atoms, and run time details are given in Table 1. For AA simulations we have adapted CHARMM36 force-field parameters given in the work of Gumbart et al. ^1^ Molecular topology files for glycan strands are provided in the Supporting Information (SI), while the topology files for the networks are available upon request.

**Table 1:**
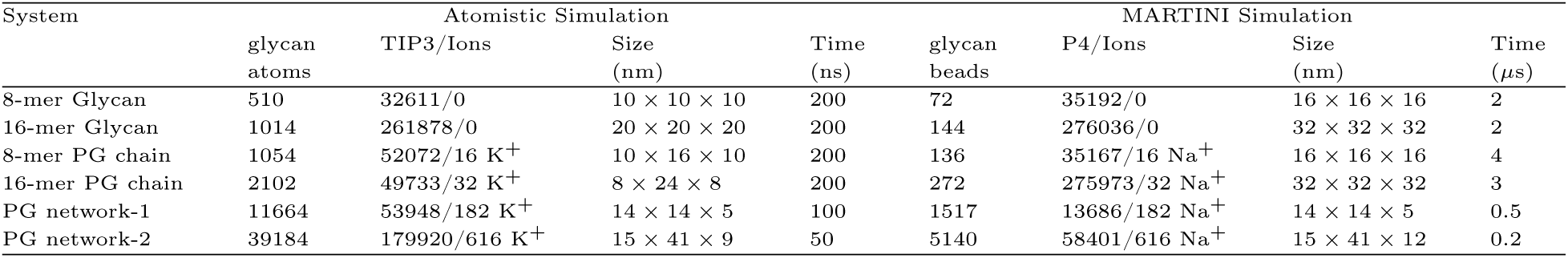
Summary of simulation systems

**Figure 2:**
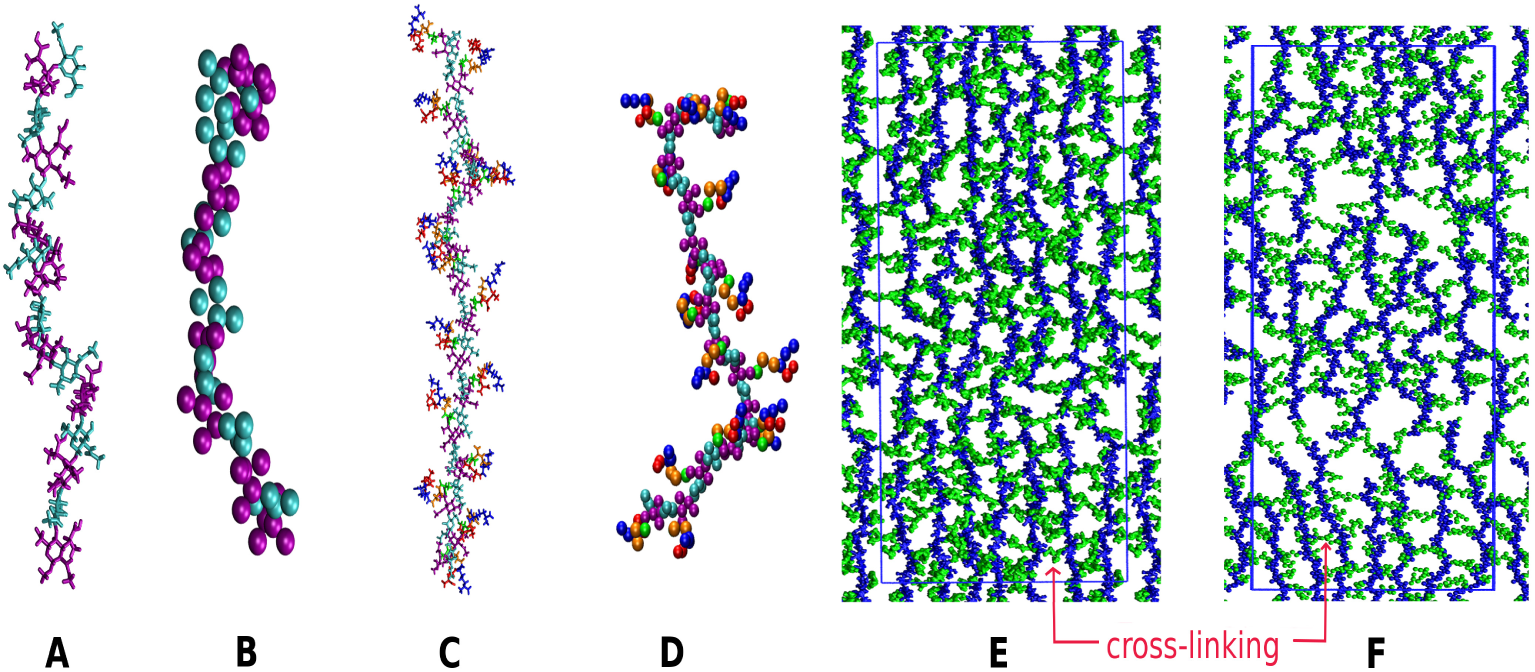
Simulated systems: (from left to right) an 8-mer glycan strand in all-atom (A) and MARTINI (B) simulations, a 16-mer peptidoglycan chain in all-atom (C) and MARTINI (D), and a network composed of 21 peptidoglycan strands in all-atom (E) and MARTINI (F) simulations. The color codes for (A)-(D) refer to NAG (cyan), NAM (purple), L-ALA (green), GLU (orange), DAP (blue) and D-ALA (red), while for the network of glycan strands the sugar backbones are represented by blue color and peptides are shown in green. The rectangular box shows the periodic boundaries in x-y plane. Solvent and ions are not shown for visual clarity.

Referring to Figure 1, a glycan strand comprises of alternating units of N-acetylglucosamine (NAG) and N-acetylmuramic acid (NAM). The glycan strand in peptidoglycan has peptide stems covalently bonded with D-lactoyl groups of the NAM residues. In the network of peptidoglycan, glycan strands are cross-linked via their peptides by bridging a bond between the DAP residue in an acceptor peptide and the penultimate D-ALA residue of a donor peptide, as illustrated in Figure 1B. It is to be noted that during the dimerization of peptides, the terminal D-ALA in the donor peptide is eliminated.

### 2.1 Atomistic Simulations

Below we describe all-atom (AA) simulation details for each of the systems given in Table 1. The atom types, mass, partial atomic charges, etc. are specified in the molecular topology files provided in the SI.

#### 2.1.1 Glycan strands in water

The systems comprised of 8-mer and 16-mer glycan strands embedded in water are simulated in an isothermal-isobaric (NPT) ensemble. A constant temperature of 310 K is maintained using a Nosé-Hoover thermostat with a time constant 1 ps. ^43,44^ An isotropic pressure coupling with a coupling constant 5 ps and compressibility 4.5 × 10^5^ bar^−1^ is imposed using the Parrinello-Rahman barostat.^45^ Verlet cut-off scheme is chosen to compute the non-bonded interactions (Lennard-Jones 6-12), which are truncated at the cut-off radius of 1.2 nm. The non-bonded van der Waals interactions are shifted with a force-switch function between 1.0 to 1.2 nm to render the forces continuous at the cut-off radius. The long ranged Coulomb interactions are computed using the Particle Mesh Ewald^46^ method with a real-space cut-off 1.2 nm. The periodic boundary condition is used in all three directions. The CHARMM TIP3P is employed as a solvent. The positions and velocities of the atoms are updated with a time step size of 2 fs using the leap-frog algorithm.

#### 2.1.2 Peptidoglycan chains in water

An 8-mer strand of glycan with short peptide of five amino acids is simulated under the conditions mentioned above. The residues GLU and terminal D-ALA have a unit negative charge, and therefore the system is neutralized by adding potassium ions. A trajectory of 200 ns is generated, and the last 100 ns trajectory is analyzed to obtain the distributions for bonded-interactions. In addition to this, the simulation is repeated for a 16-mer long peptidoglycan chain to verify the reproducibility of the bonded interactions.

#### 2.1.3 Networks of peptidogycan

A peptidoglycan network (PG network-1) is constructed by cross-linking 7 glycan strands through the peptides. The strands are initially configured with an inter-strand distance of ∼ 2 nm over a simulation patch of 14 × 14 nm. Each strand of glycan contains 13 units of disaccharides. In order to determine the extent of cross-linking between the adjacent glycan strands, a trial simulation is performed with a harmonic restraint of *f* = 400 kJ mol^−1^nm^−2^ on the heavy atoms (carbon and oxygen) in sugar rings, while the peptide residues are kept restraint-free. Based on the frequency of contacts with a cut-off distance of 0.8 nm between the free amino group of DAP residue in acceptor peptides and the carbonyl group of DALA in donor peptides, the adjacent glycan strands are cross-linked preferentially with 50% linkages.

In addition, a bigger size network of peptidoglycan (PG network-2) comprising of 21 glycan strands of varying length (on average ∼ 15 disaccharides long) is simulated for evaluating structural properties. The glycan strands and the peptides are covalently linked across the periodic boundaries, representing an infinitely large network of peptidoglycan.

### 2.2 Coarse-grained simulations

The MARTINI constructs equivalent to the AA systems just described above are simulated using GROMACS. The MARTINI bead type, mass, charge and other bonded information are given in molecular topology files in SI. The system size and duration of simulations are given in Table 1.

#### 2.2.1 Glycan strands in MARTINI framework

The MARTINI simulations are carried out on systems with glycan strands in MARTINI water. The glycan strands are 8-mer and 16-mer long in size. The thermostating of 310 K is achieved through velocity rescaling^47^ (v-rescale) with a time constant *τ*_*t*_ = 1 ps, and the pressure of 1 bar is maintained using the barostat of Parrinello-Rahman^45^ with a time constant *τ*_*p*_ = 12 ps and compressibility 3 × 10^−4^ bar^−1^. The systems are periodic in three dimensions. The van der Waals (LJ) interactions are evaluated using Verlet cut-off scheme with a cut-off distance of 1.1 nm. Equations of motion are integrated using the leap frog algorithm with a time stepping of 20 fs. The optimized parameters for bonded-interactions, namely bonds, angles and dihedrals, are provided in Tables 2-4. The simulations with the optimized parameters are extended upto 2 *µ*s. We have used MARTINI force field (version 2.2), with scaling down the energy parameter *ϵ* for LJ interactions.

**Table 2:** MARTINI parameters for bonds in glycan strand. Bead labels are according to Figure 1

| Label | Beads | Bond-length $r_o$<br>(nm) | Stiffness $k_b$<br>(kJ mol<sup>-1</sup> nm<sup>-2</sup>) |
| --- | --- | --- | --- |
| B1 | 1-2 | 0.330 | constraint |
| B2 | 1-3 | 0.300 | 8800 |
| B3 | 1-4 | 0.309 | constraint |
| B4 | 2-4 | 0.350 | 30000 |
| B5 | 5-6 | 0.331 | constraint |
| B6 | 5-7 | 0.290 | 19000 |
| B7 | 5-8 | 0.312 | constraint |
| B8 | 6-8 | 0.348 | 21000 |
| B9 | 8-9 | 0.299 | 11050 |
| B10 | 1-8 | 0.292 | 20500 |
| B11 | 5-21 | 0.294 | 20500 |

**Table 3:** MARTINI parameters for angles in glycan strand. Figure 1 can be referred for bead labels

| Label<sup>a</sup> | Beads | Angle $\theta_o$<br>(degree) | Stiffness $k_a$<br>(kJ mol<sup>-1</sup>) |
| --- | --- | --- | --- |
| A1 | 3-1-2 | 149 | 50 |
| A2 | 3-1-4 | 82 | 150 |
| A3 | 7-5-6 | 156 | 160 |
| A4 | 7-5-8 | 83 | 330 |
| A5 | 9-8-5 | 133 | 620 |
| A6 | 9-8-6 | 125 | 150 |
| A7 | 2-1-8 | 94 | 140 |
| A8 | 3-1-8 | 111 | 60 |
| A9 | 4-1-8 | 143 | 114 |
| A10 | 1-8-6 | 72 | 400 |
| A11 | 1-8-9 | 90 | 100 |
| A12 | 1-8-5 | 128 | 480 |
| A13 | 6-5-21 | 82 | 110 |
| A14 | 7-5-21 | 131 | 70 |
| A15 | 8-5-21 | 146 | 240 |
| A16 | 5-21-19 | 71 | 430 |
| A17 | 5-21-18 | 134 | 530 |

<sup>a</sup>restricted bending potential for angles A7-A17.

**Table 4:**
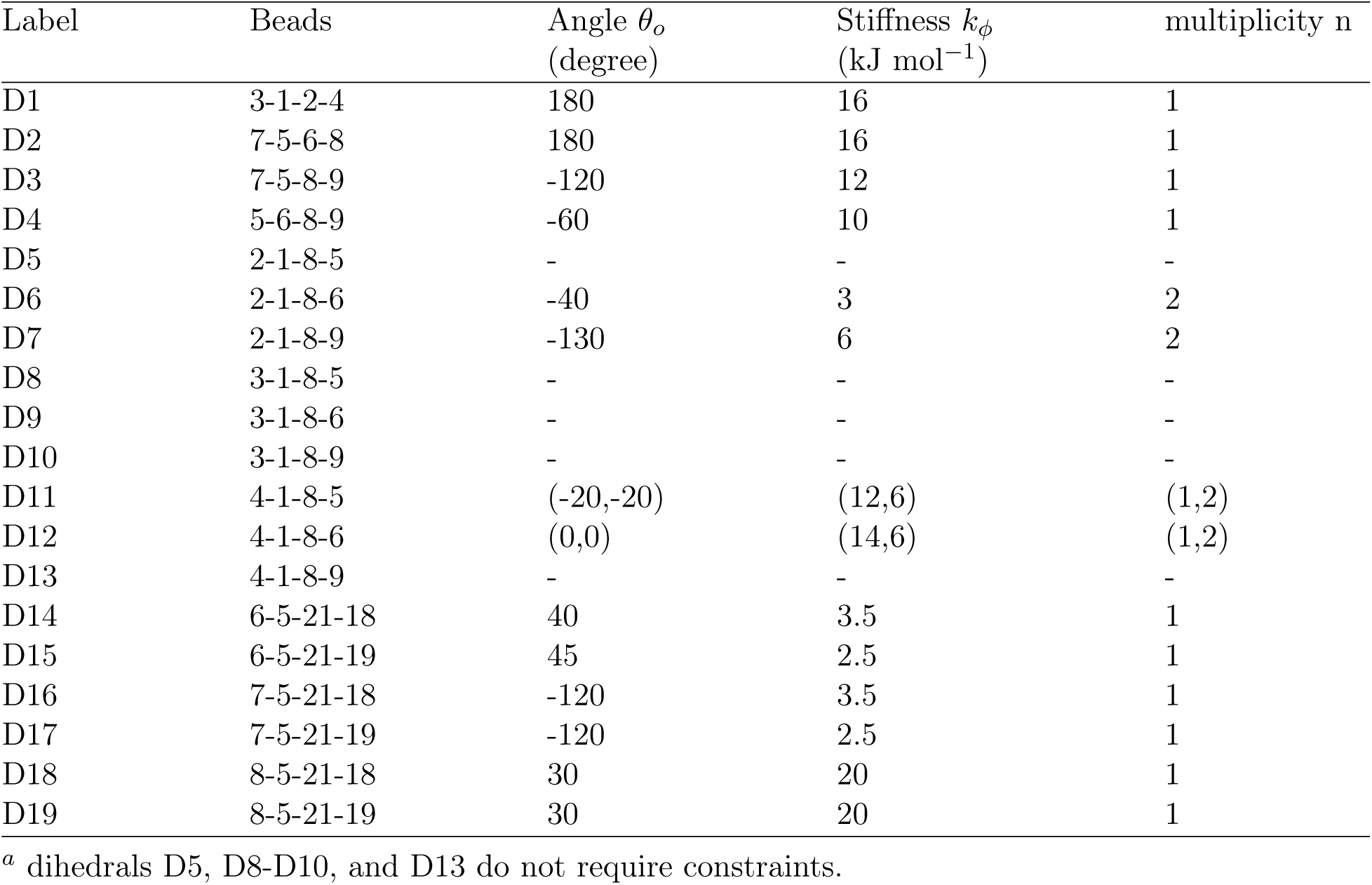
MARTINI parameters for dihedrals in glycan strand. Bead labels are according to Figure 1

| Label | Beads | Angle $\theta_o$<br>(degree) | Stiffness $k_\phi$<br>(kJ mol<sup>-1</sup>) | multiplicity n |
| --- | --- | --- | --- | --- |
| D1 | 3-1-2-4 | 180 | 16 | 1 |
| D2 | 7-5-6-8 | 180 | 16 | 1 |
| D3 | 7-5-8-9 | -120 | 12 | 1 |
| D4 | 5-6-8-9 | -60 | 10 | 1 |
| D5 | 2-1-8-5 | - | - | - |
| D6 | 2-1-8-6 | -40 | 3 | 2 |
| D7 | 2-1-8-9 | -130 | 6 | 2 |
| D8 | 3-1-8-5 | - | - | - |
| D9 | 3-1-8-6 | - | - | - |
| D10 | 3-1-8-9 | - | - | - |
| D11 | 4-1-8-5 | (-20,-20) | (12,6) | (1,2) |
| D12 | 4-1-8-6 | (0,0) | (14,6) | (1,2) |
| D13 | 4-1-8-9 | - | - | - |
| D14 | 6-5-21-18 | 40 | 3.5 | 1 |
| D15 | 6-5-21-19 | 45 | 2.5 | 1 |
| D16 | 7-5-21-18 | -120 | 3.5 | 1 |
| D17 | 7-5-21-19 | -120 | 2.5 | 1 |
| D18 | 8-5-21-18 | 30 | 20 | 1 |
| D19 | 8-5-21-19 | 30 | 20 | 1 |

<sup>a</sup> dihedrals D5, D8-D10, and D13 do not require constraints.

#### 2.2.2 Peptidoglycan chains in MARTINI framework

An 8-mer long peptidoglycan chain composed of sugars and peptides is modelled using MARTINI beads, and simulated at the conditions mentioned above. Sodium ions (MARTINI v.2.0) are added to neutralize the charges on DAP and terminal D-ALA residues. The long ranged electrostatic interactions are treated using the reaction field method ^48^ with a screening constant, *ϵ*_*r*_ = 15. The optimized parameters for bonds, angles and dihedrals are given in Tables 5-7. The simulation run time is 4 *µ*s. In order to verify the robustness of the model parameters, the simulation is repeated with a 16-mer long chain.

**Table 5:** MARTINI parameters for bonds in peptidoglycan strand. Bead labels are as per Figure 1

| Label | Beads<br>(nm) | Bond-length $r_o$<br>(kJ mol<sup>-1</sup> nm<sup>-2</sup>) | Stiffness $k_b$ |
| --- | --- | --- | --- |
| B1 | 1-2 | 0.334 | constraint |
| B2 | 1-3 | 0.301 | 10000 |
| B3 | 1-4 | 0.305 | constraint |
| B4 | 2-4 | 0.350 | 33000 |
| B5 | 5-6 | 0.330 | constraint |
| B6 | 5-7 | 0.290 | 16000 |
| B7 | 5-8 | 0.308 | constraint |
| B8 | 6-8 | 0.343 | 44000 |
| B9 | 8-9 | 0.300 | 21000 |
| B10 | 1-8 | 0.300 | 28500 |
| B11 | 5-21 | 0.288 | 32000 |
| B12 | 9-10 | 0.349 | 14800 |
| B13 | 10-11 | 0.354 | 8140 |
| B14 | 11-12 | 0.401 | 28000 |
| B15 | 12-13 | 0.315 | 8230 |
| B16 | 13-14 | 0.343 | 45000 |
| B17 | 14-15 | 0.332 | 44000 |
| B18 | 13-16 | 0.331 | constraint |
| B19 | 16-17 | 0.357 | 25700 |

**Table 6:** MARTINI parameters for angles in peptidoglycan strand. Bead labels are in accordance with Figure 1

| Label<sup>a</sup> | Beads | Angle $\theta_o$<br>(degree) | Stiffness $k_a$<br>(kJ mol<sup>-1</sup>) |
| --- | --- | --- | --- |
| A1 | 3-1-2 | 153 | 50 |
| A2 | 3-1-4 | 83 | 165 |
| A3 | 7-5-6 | 154 | 280 |
| A4 | 7-5-8 | 86 | 450 |
| A5 | 9-8-5 | 121 | 510 |
| A6 | 9-8-6 | 126 | 290 |
| A7 | 2-1-8 | 90 | 130 |
| A8 | 3-1-8 | 118 | 100 |
| A9 | 4-1-8 | 151 | 170 |
| A10 | 1-8-6 | 70 | 600 |
| A11 | 1-8-9 | 93 | 180 |
| A12 | 1-8-5 | 128 | 555 |
| A13 | 6-5-21 | 113 | 90 |
| A14 | 7-5-21 | 99 | 100 |
| A15 | 8-5-21 | 145 | 250 |
| A16 | 5-21-19 | 76 | 400 |
| A17 | 5-21-18 | 135 | 970 |
| A18 | 8-9-10 | 104 | 80 |
| A19 | 9-10-11 | 107 | 60 |
| A20 | 10-11-12 | 92 | 100 |
| A21 | 11-12-13 | 117 | 80 |
| A22 | 12-13-16 | 108 | 280 |
| A23 | 13-16-17 | 102 | 220 |
| A24 | 12-13-14 | 99 | 80 |
| A25 | 13-14-15 | 151 | 230 |
| A26 | 16-13-14 | 98 | 285 |

<sup>a</sup>restricted bending potential for angles A9-A26.

**Table 7:** MARTINI parameters for dihedrals in peptidoglycan strand. Figure 1 can be referred for bead labels

| Label<sup>a</sup> | Beads<br>(degree) | Angle $\theta_o$<br>(kJ mol<sup>-1</sup>) | Stiffness $k_\phi$ | multiplicity n |
| --- | --- | --- | --- | --- |
| D1 | 3-1-2-4 | 180 | 10 | 1 |
| D2 | 7-5-6-8 | 180 | 10 | 1 |
| D3 | 7-5-8-9 | (180,0) | (7,3) | (1,2) |
| D4 | 5-6-8-9 | (0,0) | (5,5) | (1,2) |
| D5 | 2-1-8-5 | (0,0) | (3.5,3) | (1,2) |
| D6 | 2-1-8-6 | (0,0) | (3.5,3) | (1,2) |
| D7 | 2-1-8-9 | - | - | - |
| D8 | 3-1-8-5 | - | - | - |
| D9 | 3-1-8-6 | - | - | - |
| D10 | 3-1-8-9 | (0,0) | (8,2) | (1,2) |
| D11 | 4-1-8-5 | - | - | - |
| D12 | 4-1-8-6 | (0,0) | (7,3) | (1,2) |
| D13 | 4-1-8-9 | (180,180) | (2,3) | (2,3) |
| D14 | 6-5-21-18 | (180,0) | (6,4) | (1,2) |
| D15 | 6-5-21-19 | - | - | - |
| D16 | 7-5-21-18 | - | - | - |
| D17 | 7-5-21-19 | - | - | - |
| D18 | 8-5-21-18 | (0,0) | (8,2) | (1,2) |
| D19 | 8-5-21-19 | (0,0) | (7,3) | (1,2) |
| D20 | 8-9-10-11 | 60 | 3 | 2 |
| D21 | 9-10-11-12 | 140 | 3 | 2 |
| D22 | 10-11-12-13 | (180,0) | (5,5) | (1,2) |
| D23 | 11-12-13-16 | 180 | 3.5 | 1 |
| D24 | 12-13-16-17 | (180,0) | (7,5) | (1,2) |
| D25 | 11-12-13-14 | (180,0) | (5,5) | (1,2) |
| D26 | 12-13-14-15 | 0 | 5 | 1 |
| D27 | 16-13-14-15 | (180,0) | (5.2,3.8) | (1,2) |
| D28 | 17-16-13-14 | 180 | 6 | 1 |

<sup>a</sup> dihedrals D7-D9, D11, and D15-D17 do not require constraints.

#### 2.2.3 Peptidoglycan networks in MARTINI framework

A CG network of peptidoglycan (PG network-1) is prepared by assembling and cross-linking the glycan strands, 7 in number, through the peptides. With an inter-strand spacing of ∼ 2 nm, the length of each glycan strand is 13 disaccharides. In order to connect the peptides to form preferentially dimeric linkages, like in AA simulations, a trial MARTINI simulation is carried out with a harmonic restraint of 150 kJ mol^−1^ nm^−2^ on one of the beads in each sugar rings (e.g. beads 2,6,19 etc. in Figure 1). The frequency of contacts between the DAP residue in acceptor peptides and penultimate D-ALA in donor peptides (e.g. beads forming a cross-link in Figure 1) within a cut-off distance of 0.8 nm determines the potential pairs of peptides to be covalently linked. The percentage cross-linking is restricted to ∼ 50%.

A larger network of peptidoglycan (PG network-2) having size ∼ 15 × 41 nm is also constructed using 21 CG glycan strands. The network serves as a more realistic cell wall model, as the glycan strands and peptides are covalently bonded across periodic boundaries. This larger simulation patch is employed for free energy calculations as well as for estimating the mechanical properties, namely area compressibility and bending modulus of the peptidoglycan. The surface-tension parameter in GROMACS is employed for anisotropic pressure coupling to set a desired tension in the MARTINI network.

For assessing the free energy barriers for small molecules, the umbrella sampling simulations were carried out for thymol insertion though a monolayer of peptidoglycan. Thymol was parametrized using CGenFF^49^ to generate its AA force fields, while an automated parametrization scheme introduced by Bereau and Kremer ^50^ was used for coarse-graining of thymol. It should be noted that the van der Waals interactions between the peptidoglycan and thymol beads in MARTINI simulations are scaled according to equation (1) using the parameter *α* = 0.7, which was used to scale the interaction between the peptidoglycan beads. The interactions of thymol and peptidoglycan with water and ions are unscaled. The CG mapping scheme and the MARTINI bead types for thymol (Figure S1), as well as the histograms for umbrella sampling biasing potential (Figures S2-S4) are given in SI.

For the smaller patch of peptidoglycan (PG network-1) the oxygen atoms in sugar backbones in AA model are harmonically restrained by a force constant *f*_*z*_ = 500 kJ mol^−1^ nm^−2^, while a weak restraint is imposed on the CG beads (2,6,19 etc. in Figure 1) containing the oxygen in sugar rings in MARTINI simulations to keep the peptidoglycan sheet more or less planar. The umbrella sampling simulations are performed on a reaction coordinate, which is the distance between the center of mass of the peptidoglycan sheet and the center of mass of thymol along the z-direction-normal to the sheet of peptidoglycan. Steered molecular dynamics simulations are employed to generate initial configurations for the umbrella sampling simulations,^51^ and the sampling windows are created at a spacing of 0.1 nm. The thymol molecule is restrained using a harmonic potential with a force constant of 1000 kJ mol^−1^ nm^−2^. Each umbrella window is simulated for 100 ns and 500 ns in AA and MARTINI simulations, respectively. The free energy profile is computed by using the weighted histogram analysis method (WHAM),^52^ which is implemented in GROMACS. The MARTINI beads are evolved with a time step size of 10 fs in all simulations on the peptidoglycan networks.

### 2.3 Model Parameters

As mentioned earlier, the MARTINI force field (version 2.2) is adapted for non-bonded interactions. The original MARTINI parameters, namely energy *ϵ* and size *σ*, lead to an aggregation of the CG beads for polysaccharides in MARTINI simulations.^36^ It has been suggested to increase the repulsion between CG beads by reducing the energy parameter *ϵ* to remedy the issue of bead aggregation. In order to circumvent the aggregation of CG beads observed in our MARTINI simulations (Figure S5 in SI), we increased the bead-bead repulsion by scaling down *ϵ* with a parameter (*α*) using,

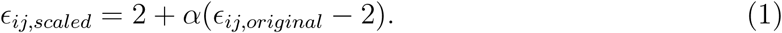

We varied *α* in the range 0.9 to 0.7, and found that the scaling factor *α* = 0.7 is sufficient to avoid bead aggregation and that it reproduces the chain end-to-end distance close to the end-to-end distance obtained in all-atom simulations (Table 8). By increasing the bead-bead repulsion in the MARTINI force field, we have introduced new bead types, which are distinguished from original MARTINI beads using the letter R. For example, a bead P1 in the original MARTINI force field is now recognized as RP1 with the scaled values of *ϵ*. The interactions of R beads with water (P4) and sodium ions (Q_*d*_) are unscaled.

**Table 8:** Summary on properties of peptidoglycan

| Property | MARTINI simulation | AA (this work) | Literature |
| --- | --- | --- | --- |
| End-to-end distance | 3.8 nm (8-mer)<br>7.5 nm (16-mer) | 4.1 (8-mer)<br>8.1 (16-mer) | -<br>- |
| Peptides orientation | $\sim 90^\circ$ | $\sim 90^\circ$ | $80 - 100^\circ$,<sup>18</sup> $75 - 105^\circ$<sup>19</sup> |
| Thickness | 4 – 4.5 nm | 4 – 4.5 nm | 4 nm,<sup>10</sup> 3.4 – 3.9 nm<sup>1</sup> |
| Area per disaccharide | 2 nm<sup>2</sup> | 2 nm<sup>2</sup> | 2.5 nm<sup>2</sup>,<sup>55</sup> 2.6 – 3.1 nm<sup>2</sup><sup>1</sup> |
| Area compressibility | 20 – 100 mN/m | - | 29 – 500 mN/m<sup>21</sup> |
| Bending modulus | $\sim 1$ k<sub>B</sub>T | $\sim 1$ k<sub>B</sub>T | - |
| Insertion free energy<br>for thymol | $\sim -1$ kJ/mol | $\sim -1$ kJ/mol | - |

The bonded interactions, namely bond-stretching (two body) and angle-bending (three body), are modelled using harmonic springs and cosine potentials, respectively,

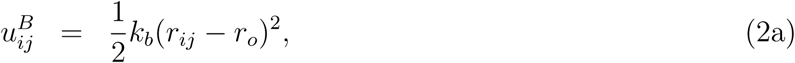

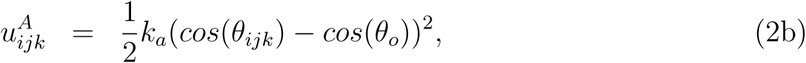

where *k*_*b*_ and *k*_*a*_ represent the bond and angle-bending stiffness constants, while the equilibrium bond lengths and angles are denoted by *r*_*o*_ and *θ*_*o*_, respectively. In order to avoid numerical instability arising from torsion angle calculations, some of the angles are maintained at their equilibrium values using the restricted bending potential,^53^

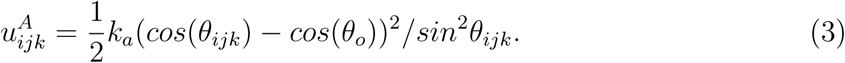

The torsional angle among quadruple of beads *i* − *j* − *k* − *l* is controlled using cosine function,

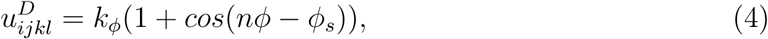

with *ϕ* being the torsion angle between the planes formed by the triplets *i* − *j* − *k* and *j* − *k* − *l*. The stiffness *k*_*ϕ*_, multiplicity *n* and dihedral angle *ϕ*_*s*_ are the model parameters.^54^

The optimized parameters for bonded interactions within the glycan strands are given in Table 2 for bonds, in Table 3 for angles, and in Table 4 for dihedrals. For the peptidoglycan chains the optimized parameters are tabulated in Tables 5-7. When two glycan strands are cross-linked via their peptides, there are additional bonded interactions involving a bond that bridges the peptides and the angles surrounding this bond. The equilibrium bond length at cross-links is 0.35 nm and is maintained with a harmonic potential of strength 1100 kJ mol^−1^ nm^−2^.

## 3 Results and Discussion

In this section we explain the procedure for optimizing model parameters, the mapping scheme, selection of MARTINI bead types and validation of the CG models.

### 3.1 Model Development

The model parameters for bonded interactions, namely the bond-stiffness *k*_*b*_, equilibrium bond length *r*_*o*_, angle-bending stiffness *k*_*a*_, equilibrium angle *θ*_*o*_, and dihedral parameters, namely *k*_*ϕ*_, *ϕ*_*s*_ and multiplicity *n*, are optimized by mapping the bond, angle and dihedral distributions from MARTINI simulations with the corresponding reference distributions obtained in atomistic simulations. Toward this end, virtual CG trajectories are derived from AA trajectories from the center of mass of groups of atoms forming the CG beads. Figure 1 shows a mapping scheme employed during the coarse-graining of peptidoglycan. The coarse-grained beads are assigned appropriate MARTINI bead types based on the functionality (polarity and hydrogen bonding capability) of the underlying atoms within the beads. The particle type ‘S’ is ascribed to smaller size beads in the ring structures of sugar units to mimic a relatively smaller (3:1 or 2:1) mapping compared to the regular 4:1 mapping scheme of MARTINI. Within the MARTINI framework, the polar groups of atoms are labelled with ‘P’ type, while the bead type ‘N’ is used for groups which are partly polar and partly apolar. The charged beads are denoted by ‘Q’. The apolar beads in peptides are regarded as ‘AC’ for their intra-interactions with ‘Q’ type particles of the peptides.

To begin an iterative process of MARTINI parametrization, the centers of histograms for bonds and angles from AA simulations of an an 8-mer glycan strand are used as an initial set of equilibrium bond lengths and angles in MARTINI simulations. These values are updated after every 500 ns of MARTINI simulation according to,

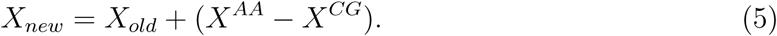

Here *X* is the equilibrium bond length (angle) for bond (angle) distributions, while *X*^*AA*^ and *X*^*CG*^ are centered at the corresponding distributions in AA and MARTINI simulations, respectively. The strengths of bond-stretching (*k*_*b*_) and angle-bending (*k*_*a*_) are updated according to the heights of the distributions, which are *Y* ^*AA*^ in AA and *Y* ^*CG*^ in the MARTINI simulations, using

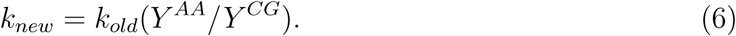

The stiffness constant *k* represents *k*_*b*_ for the bond distributions and *k*_*a*_ for the angle distribution. The iterative procedure terminates when the bond length in MARTINI simulation converges to its AA counterpart within a tolerance of 0.01 nm and the peak of the distribution lies within 1%. Similarly, the angles are said to be converged when the deviation from their AA target is within 2^*o*^ and the peak differences are within 1%. The procedure took 3 iterative CG simulations to converge bonds and angles before the dihedrals were set in, and with 8 additional CG simulations the bonded distributions, including 28 dihedrals, were converged. For the bonds showing multi-modal distributions, for instance the bonds labelled by B4 and B8 in Figure 3, we have parametrized the stiffness constant to realize wider CG distributions spanning the multiple peaks observed in the AA distributions. The heights of CG distributions are kept within 2% of the average in the peaks of bi-modal distributions. The dihedral parameters, namely the stiffness constant *k*_*ϕ*_, angle *ϕ*_*s*_ and multiplicity *n*, are estimated by considering the number and locations of the peaks in the AA target distributions for the dihedrals. The procedure for parameterization is illustrated in an algorithmic form in Figure S6 of SI.

**Figure 3:**
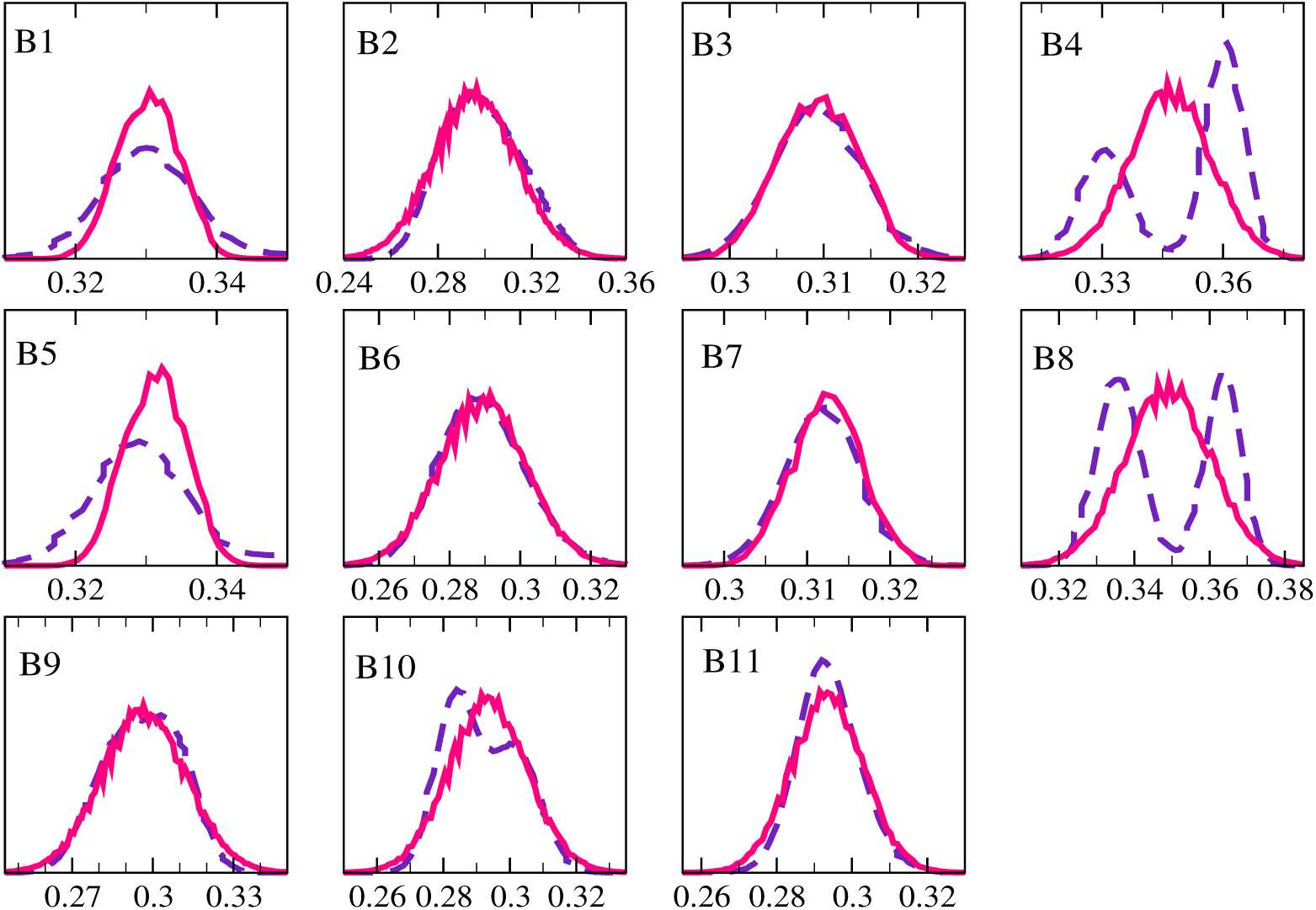
Mapping of bond distributions in MARTINI simulations with AA target dis-tributions for an 8-mer glycan strand. Symbols represent AA (−−) and MARTINI (−) distributions. The bond lengths are in nm. Table 2 can be referred for bond labels and their corresponding bond parameters.

#### 3.1.1 Modeling of Glycan Strand

The distributions for bonds, angles and dihedrals are computed from a 2 *µ*s long trajectory in a MARTINI simulation of the glycan chain comprising of 8 disaccharides. The last 1 *µ*s trajectory is used for averaging. Figure 3 shows an excellent agreement of the bond distributions of the MARTINI simulation with the AA reference distributions. The bond lengths are in the range of 0.29-0.35 nm. The bonds with stiffness constant 50000 kJ mol^−1^ nm^2^ are replaced by constraints. The distributions for angles in Figure 4 are also in consonance with their AA counterparts. The mean angles have values over a range of 70-160°. The dihedral distributions that have a single or prominent peak in AA simulations are very well captured by the MARTINI parameters for dihedrals as evidenced in Figure 5. Since the coarse-graining does not retain asymmetrical distributions of atoms that form CG clusters, mapping of dihedrals that have multiple peaks is not always captured. However for the glycan strands in question, multi-modal peaks are also mapped out satisfactorily, as apparent in Figure 5. The dihedrals having bi-modal peaks exhibit two conformers, and such dihedrals are nearly 50% in number. To test the robustness of the MARTINI parameters, we have simulated a 16-mer long glycan strand using the same model parameters obtained for the 8-mer strand. The bonded distributions for the 16-mer strand are given in SI (Figures S7-S9), and their comparison with AA data confirms the validity of the optimized parameters.

**Figure 4:**
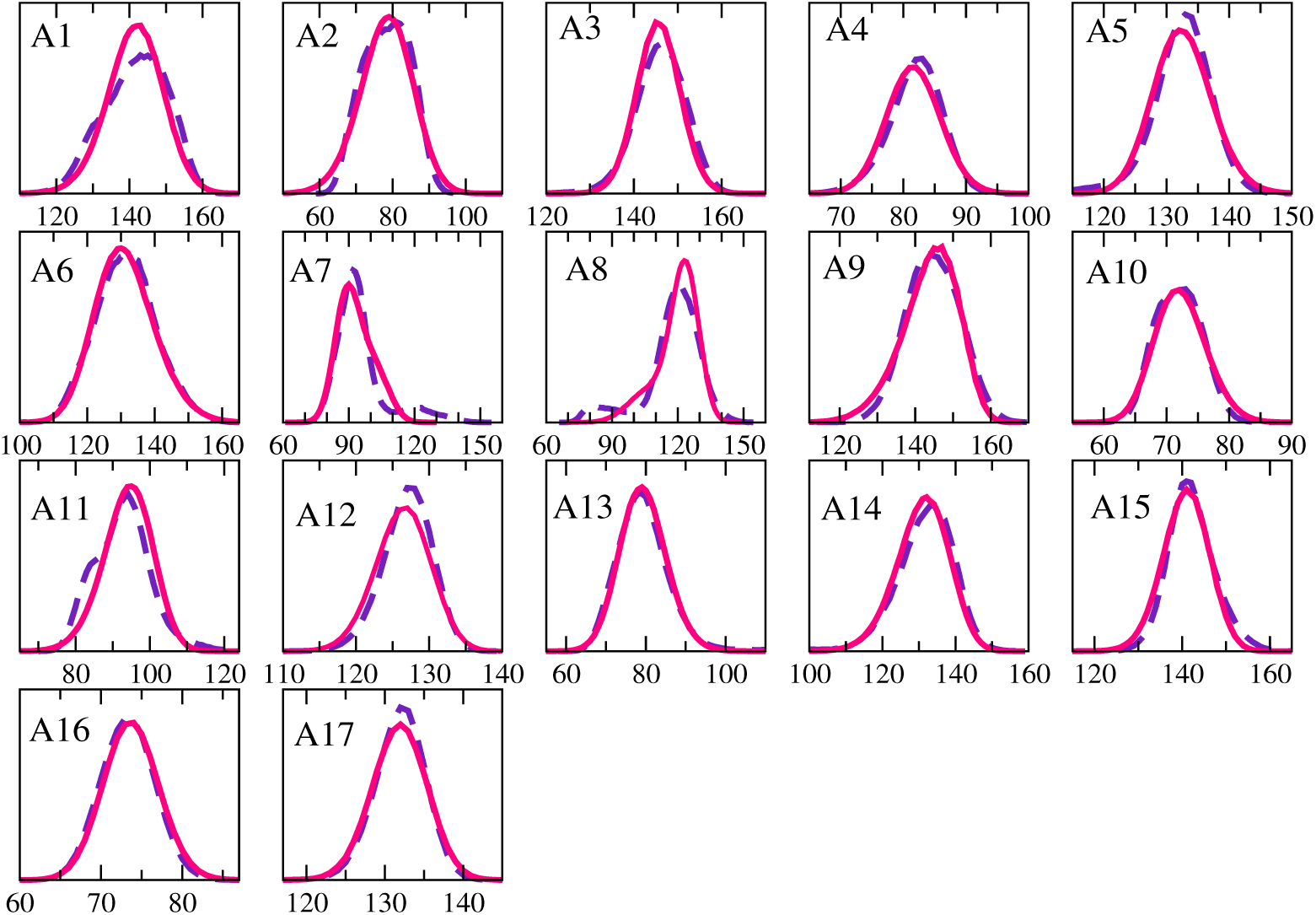
Mapping of angle distributions in MARTINI simulations with corresponding AA target distributions for an 8-mer glycan strand. Symbols represent AA (−−) and MARTINI (−) distributions. The angles are specified in degrees. The labels and angle parameters are mentioned in Table 3.

**Figure 5:**
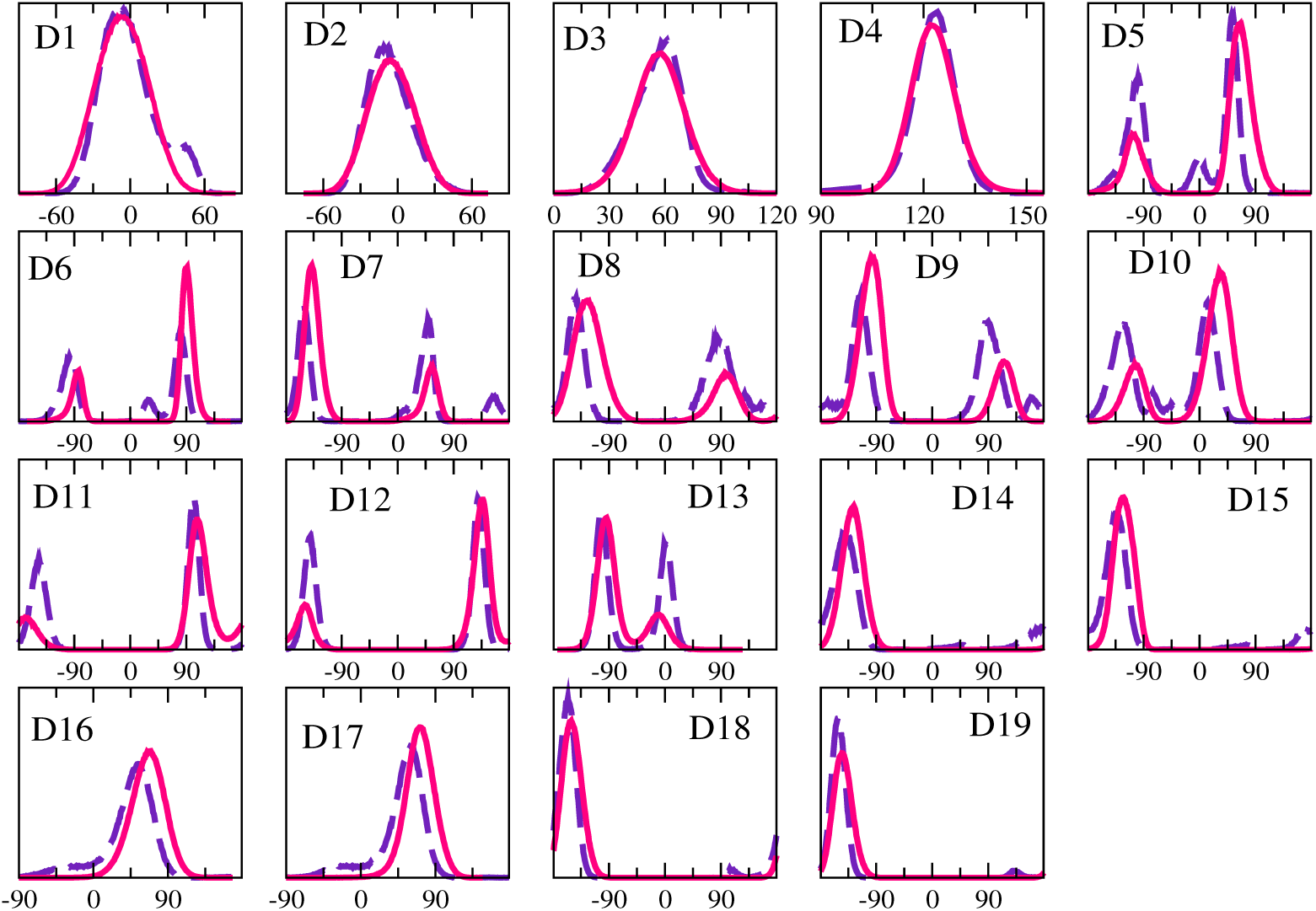
Mapping of torsional angle distributions for MARTINI simulations with corresponding AA target distributions for an 8-mer glycan strand. Symbols represent AA (−−) and MARTINI (−) distributions. The angles are specified in degrees. The labels are in accordance with Table 4.

#### 3.1.2 Modeling of Peptidoglycan

An oligomeric strand having 8 repeating units of disaccharides, with a penta-muropeptide attached to each NAM residue, is simulated using both AA force fields and a MARTINI model. A long trajectory of 2 *µ*s in MARTINI simulation is analyzed for the bonded distributions. Figures 6, 7, and 8 compare the histograms for bonds, angles and dihedrals, respectively, obtained in MARTINI simulations with those obtained using AA simulations. The matching of the profiles for bonds and angles is excellent. Most of the dihedrals are very well mapped out onto their corresponding AA target distributions. The bonded parameters given in Tables 5-7 reproduce the bonded distributions for larger lengths of the peptidoglycan chains, as evidenced in the bonded distributions for a 16-mer peptidoglycan strand in SI (Figures S10-S12).

**Figure 6:**
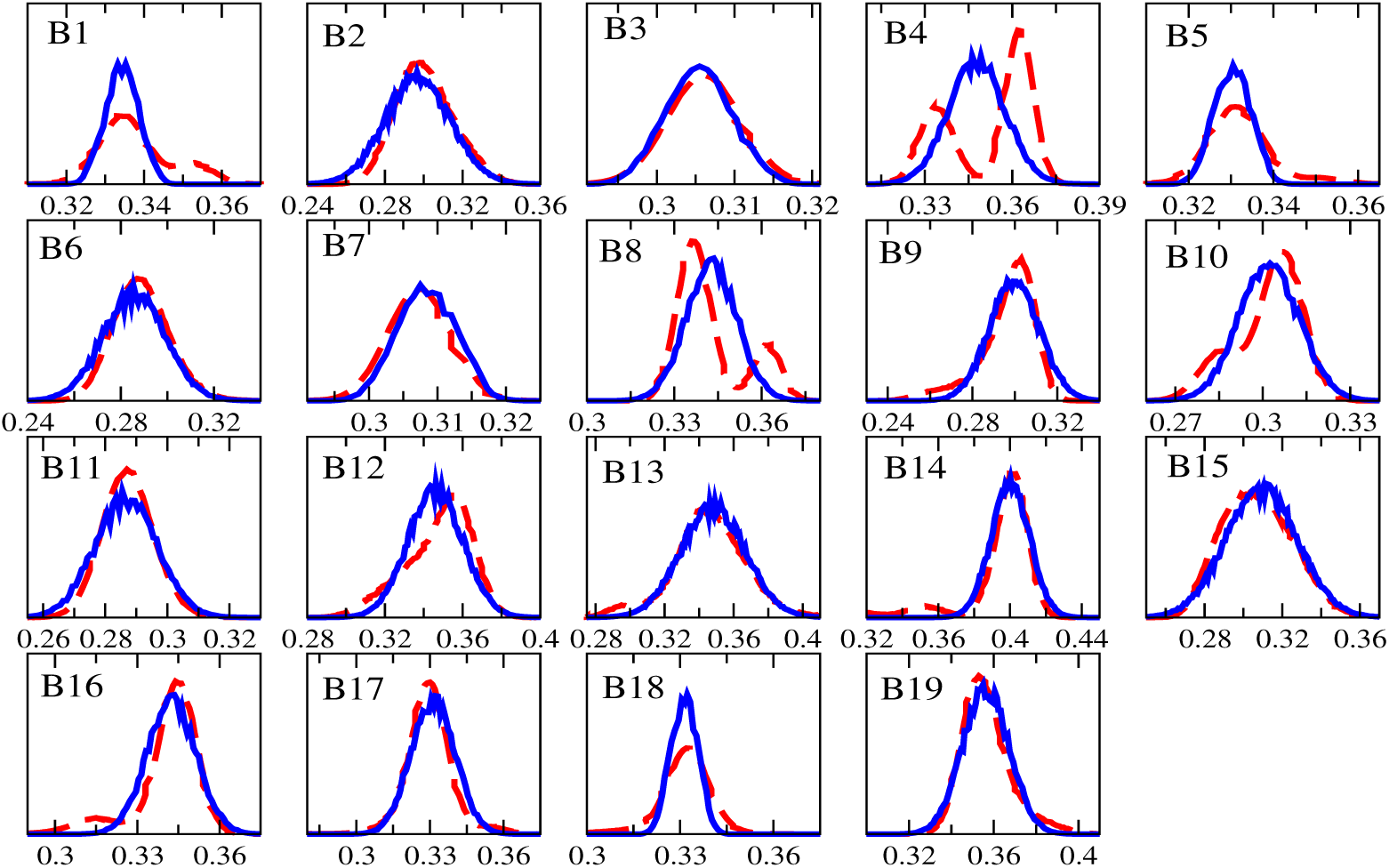
Mapping of bond length distributions for MARTINI simulations with corresponding AA target distributions for an 8-mer peptidoglycan strand. Symbol (−−) indicates AA distributions, while solid lines (−) represent MARTINI distributions. The bond lengths are in nm. The labels are in accordance with Table 5.

**Figure 7:**
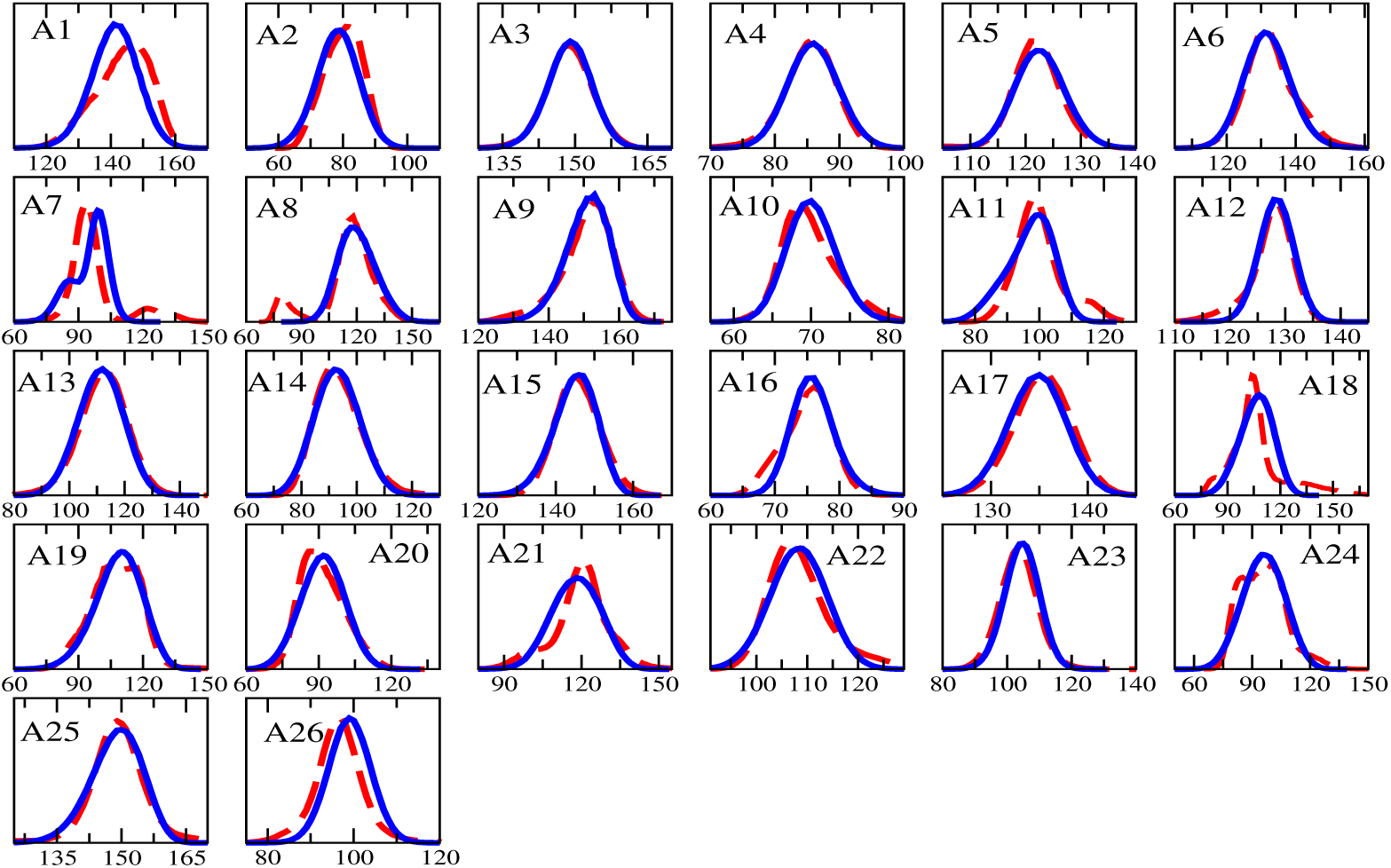
Mapping the angle distributions from MARTINI simulations with reference AA distributions for an 8-mer peptidoglycan strand. Symbols represent AA (−−) and MARTINI (−) distributions. The angles are specified in degrees. The labels and angle parameters are mentioned in Table 6.

**Figure 8:**
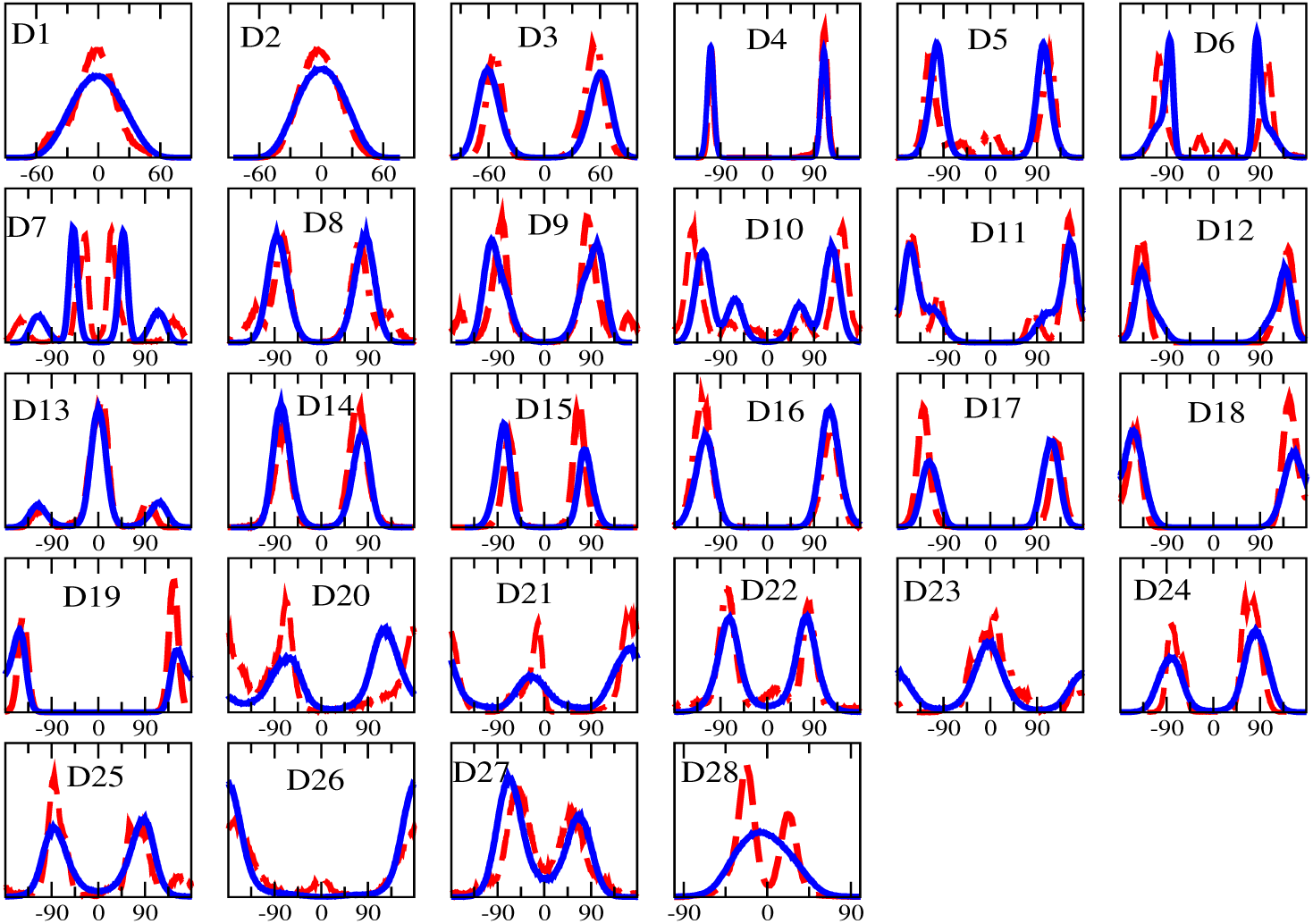
Mapping of torsional angle distributions from MARTINI simulations with reference AA distributions for an 8-mer peptidoglycan chain. Symbols represent AA (−−) and MARTINI (−) distributions. The angles are specified in degrees. Table 7 can be referred for labels and dihedral parameters.

### 3.2 Model Validation

#### 3.2.1 Glycan chain end-to-end distance

Figure 9A compares the histograms of end-to-end distance (*R*_*e*_) for the 16-mer glycan strand simulated using MARTINI and AA models. The overlap between the histograms is significantly high. The end-to-end distance for the 16-mer CG glycan strand is 7.5 ± 0.004 nm, and this is in good agreement with the end-to-end distance obtained from AA simulation of the 16-mer chain.

**Figure 9:**
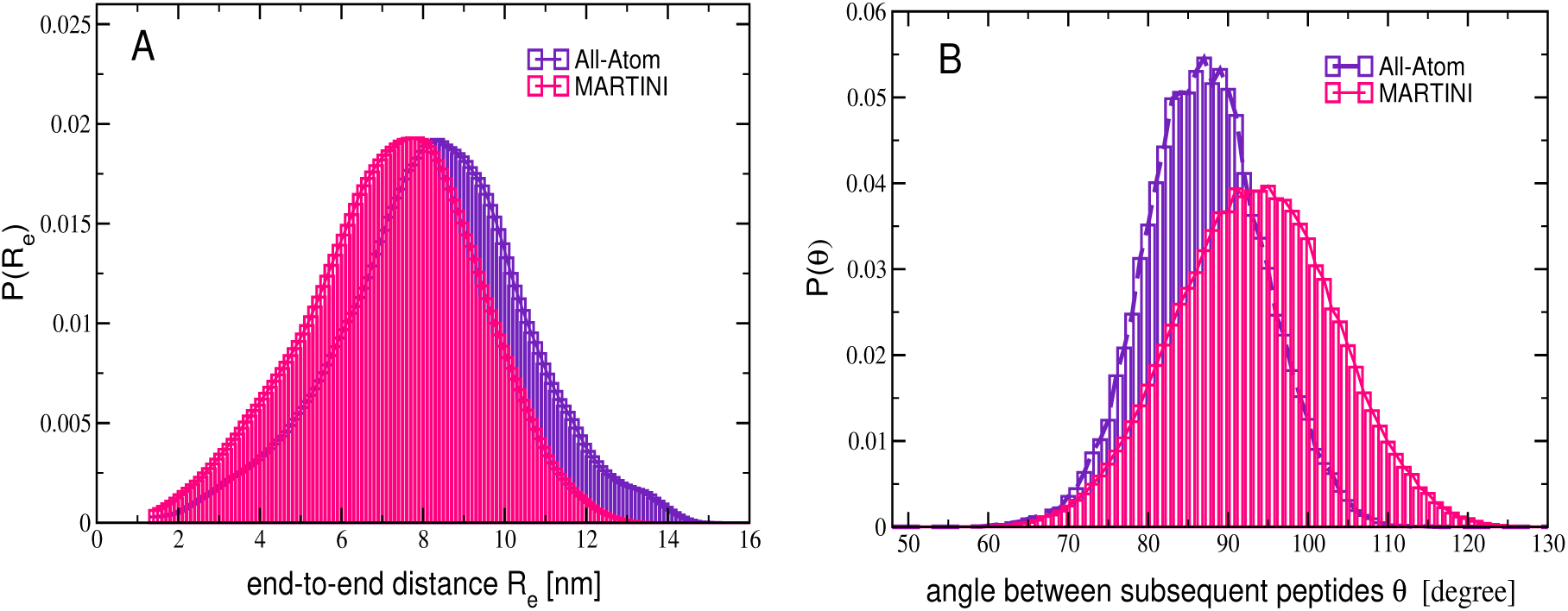
(A) Probability distributions of end-to-end distance *R*_*e*_ for the 16-mer glycan strand in AA (maroon) and MARTINI simulations (orange). (B) Probability distributions for the native angle between subsequent peptides along the peptidoglycan chain of 16-mer in AA (maroon) and MARTINI simulations (orange).

#### 3.2.2 Native periodicity of peptides

NMR spectroscopy study^20^ suggests that the successive peptides along a glycan strand are spaced ∼ 120° apart, indicating a three-fold periodicity in the peptide orientation. However, the computer simulations by Leps et al. ^18^ revealed that in energetically favorable conformations the consecutive peptides orient with a spacing of 80 − 100°, and this is further supported by other in-silico investigation reports.^1,19^ With a four-fold symmetry of 90° between the subsequent muropeptides along the glycan strand, half of the peptides would lie in the plane of glycan strands, while the remaining half would protrude out in a direction perpendicular to the plane of glycan strands. Consequently, only 50% of the peptides are accessible to form covalent bonds to cross-links with neighbouring glycan strands, supporting experimental evidence of 40-50% cross-linking.^11^ Figure 9B depicts an equilibrium angle between consecutive peptides along the sugar chain for the AA and MARTINI models. The propagation angle is found to be 93.9 ± 0.02° for MARTINI and 87.5 ± 0.03° for AA model. The distribution for the angle between consecutive peptides is broader for the MARTINI simulations due to soft coarse-grained beads. The observation of a four-fold symmetry in orientation of muropeptides is consistent with the literature.^1,18,19^

#### 3.2.3 Density distribution

Figure 10 indicates the mass density of the peptidoglycan normal to the plane of the peptidoglycan network composed of 21 glycan strands. The density profile for MARTINI model is in good agreement with that of AA model of peptidoglycan. Since total mass of the peptidoglycan is conserved between AA and MARTINI models, a small difference in peak density values is attributed to the differences in their equilibrium lateral areas, which differ only by less than 5% relative to the lateral area in the AA simulation. With a criterion of 50% of the peak density, the thickness of the peptidoglycan monolayer is ∼ 2.5 − 3.0 nm. However the thickness ∼ 4 − 4.5 nm with the 10% criterion of the peak density agrees well with other simulation literature^1^ as well as experimental tomograms.^10^

**Figure 10:**
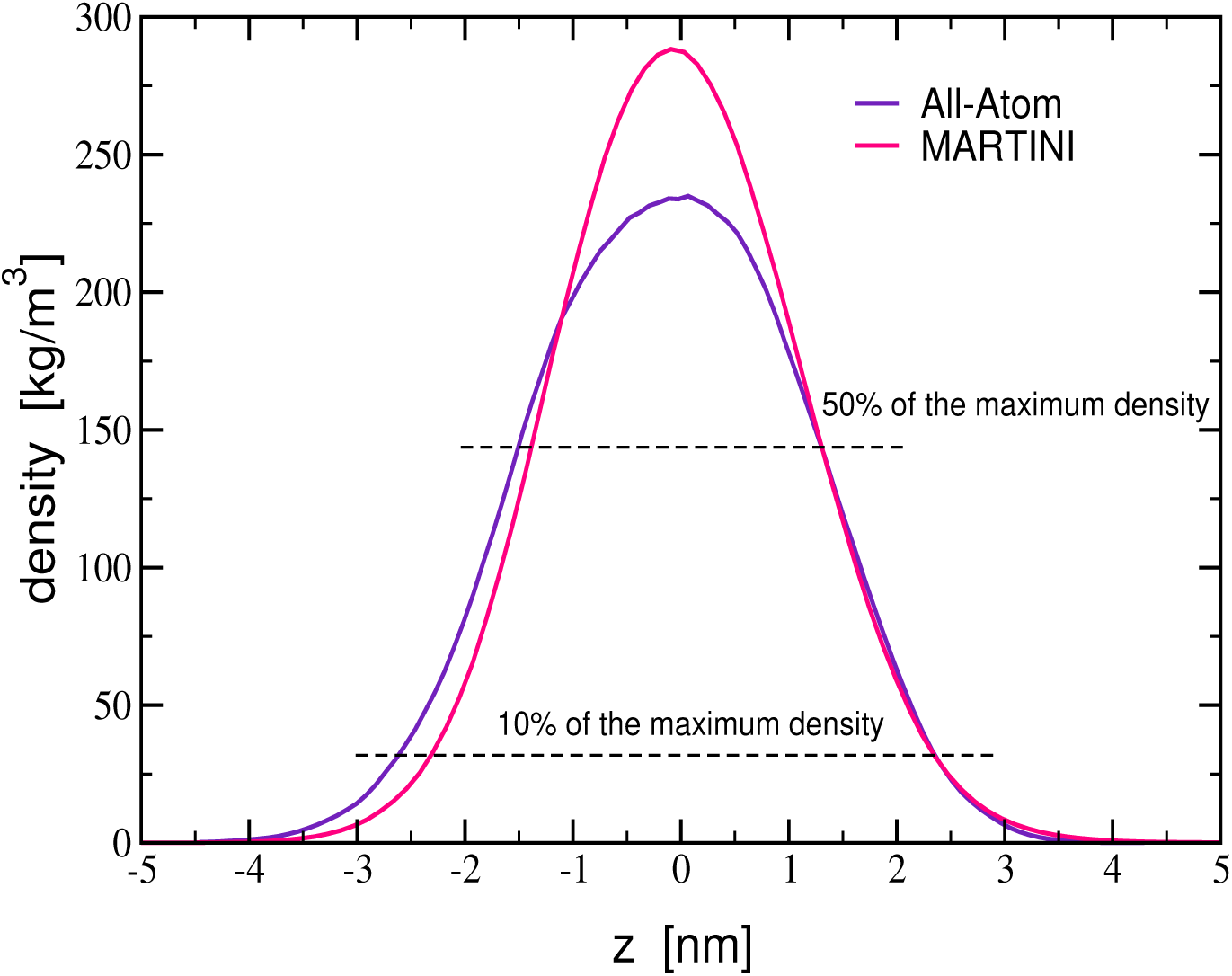
Mass density distribution of peptidoglycan in direction (z) normal to the plane of the peptidoglycan network containing 21 strands. The symbols are AA model (maroon), and MARTINI model (red). The dash lines at 10% as well as 50% of the peak density serve to estimate the thickness of the peptidoglycan layer.

The mean area per disaccharide from MARTINI simulations at ambient temperature and pressure is ∼ 2 nm^2^, which is consistent the range 2.6-3.1 nm^2^ reported in other atomistic simulations.^1^ An experimental mean area per disaccharide was estimated to be ∼ 2.5 nm^2^ using the number of DAP residues the surface area of an Escherichia coli. ^55^ This further substantiates the coarse-grained model of peptidoglycan. The area per disaccharide increases with tension in the membrane, as delineated in the next section.

#### 3.2.4 Area stretch modulus

The monolayer of peptidoglycan comprised of 21 glycan strands (Figure 2F, PG network-2) is subjected to lateral stresses, and its response in areal expansion is monitored. The surface tension is maintained at a desired value, while the pressure (*P*_*zz*_) normal to the glycan network is set to 1.0 bar. The surface tension value is varied in a range 0-50 mN/m. The surface tension (*γ*) is calculated from the lateral (*P*_*xx*_ and *P*_*yy*_) and normal (*P*_*zz*_) components of pressure tensor using, ^56^

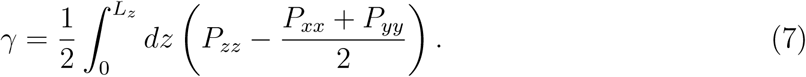

The simulation box length in z-direction is *L*_*z*_. Figure 11A shows the variation of the surface tension with the fractional change in lateral area in the plane of peptidoglycan. The lateral area increases with applied tension (Figure S13 in SI). The area compressibility (K_A_) of the network is evaluated using,

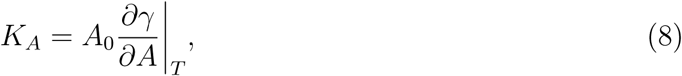

where *A*_0_ is the equilibrium area at vanishingly zero tension. The area compressibility is as low as 20 mN/m for 50% area change, and steeply increases to 100 mN/m for ∼ 100% expansion, exhibiting a strain stiffening behavior. Since the results are in good agreement with the AA simulations,^21^ the proposed CG model of peptidoglycan can be reliably used for *in-silico* investigations of bacterial cell walls under tension as high as 30 mN/m.

**Figure 11:**
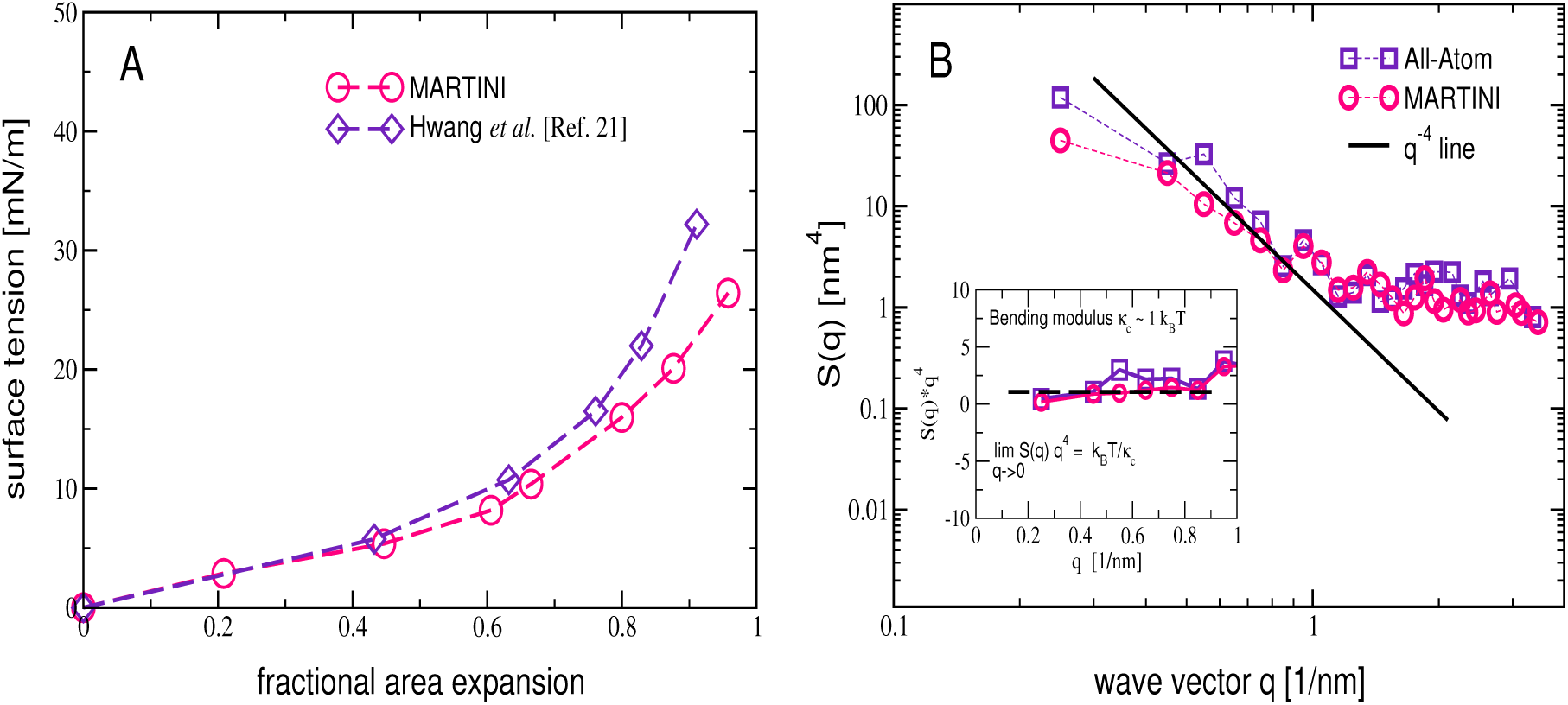
(A) Response in area expansion with tension in the MARTINI network of peptidoglycan (PG network-2), and the data represented by ⋄ are extracted from a graph from Hwang et al. ^21^ using xyscan. (B) Structure factor for height fluctuations in AA (∇) and MARTINI (Δ) representations of PG network-2, and the *q*^−4^ line is a guide to an eye. The inset shows *S*(*q*) data scaled with *q*^4^, and a horizontal dash line indicating the limiting value *k*_*B*_*T/κ*_*c*_.

#### 3.2.5 Bending modulus

We have also computed the bending modulus of the model membranes of peptidoglycan using the Helfrich analysis of the height fluctuations. ^57^ A surface is constructed through the peptidoglycan, and its thermal fluctuations are analysed in the Fourier domain. Referring to Figure 11B, the static structure factor corresponding to the height fluctuations of the peptidoglycan surface is computed using Fourier Transforms, 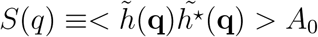, over the surface area *A*_0_ for a tensionless membrane. ^58–60^ The wave vector **q** = 2*π*(*n*_*x*_*/L*_*x*_, *n*_*y*_*/L*_*y*_), with integer numbers *n*_*x*_ and *n*_*y*_, where the linear dimensions in two orthogonal directions are *L*_*x*_ and *L*_*y*_. In the low **q** limit, the structure factor for a thermally fluctuating and tensionless two-dimensional surface follows

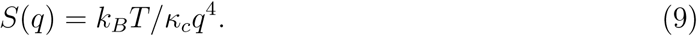

The bending modulus (*κ*_*c*_) is deduced by fitting the structure factor data to equation (9). As evident in Figure 11B the structure factor follows *q*^4^ scaling and the data for the MARTINI network of peptidoglycan are in excellent agreement with the structure factor obtained using the AA model. An inset in Figure 11B clearly indicates that the structure factor data obey a limiting behavior with an intercept k_B_T*/κ*_*c*_ ∼ 1. The low value of *κ*_*c*_ ∼ 1 k_B_T implies that a monolayer of peptidoglycan has a smaller bending modulus when compared with lipid membranes, which possess the bending modulus of typically tens of k_B_T depending on whether the lipid membrane is in gel or liquid-crystalline phase. With *κ*_*c*_ ∼ 1 k_B_T and K_a_ ∼ 10 mN/m for the tensionless network of peptidoglycan, the mechanical thickness of the network can be obtained using the elastic ratio^61^ 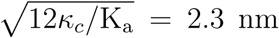. This is in good agreement with the thickness of the monolayer of the peptidoglycan discussed in the preceding section.

#### 3.2.6 Potential of Mean Force Calculations

In order to study interactions of the model peptidoglycan with small molecules, we evaluated the potential of mean force (PMF) between a thymol molecule and the peptidoglycan networks described earlier. Prior to quantifying the free energy landscapes by umbrella sampling simulations, we carried out restraint-free simulations. Figures 12A and 12B depict a translocation event observed in AA and MARTINI restraint-free simulations, respectively. The translocation of thymol through peptidoglycan occurs rapidly over a time scale of 2 ns, indicating that peptidoglycan does not offer any significant barrier for thymol. This free permeation of thymol through the peptidoglycan is further vindicated by Figure 12C, which clearly indicates the absence of any significant barrier in the PMF profiles. The densities of glycan, peptides and water are also indicated in Figure 12C to precisely mark the locations of these species.

**Figure 12:**
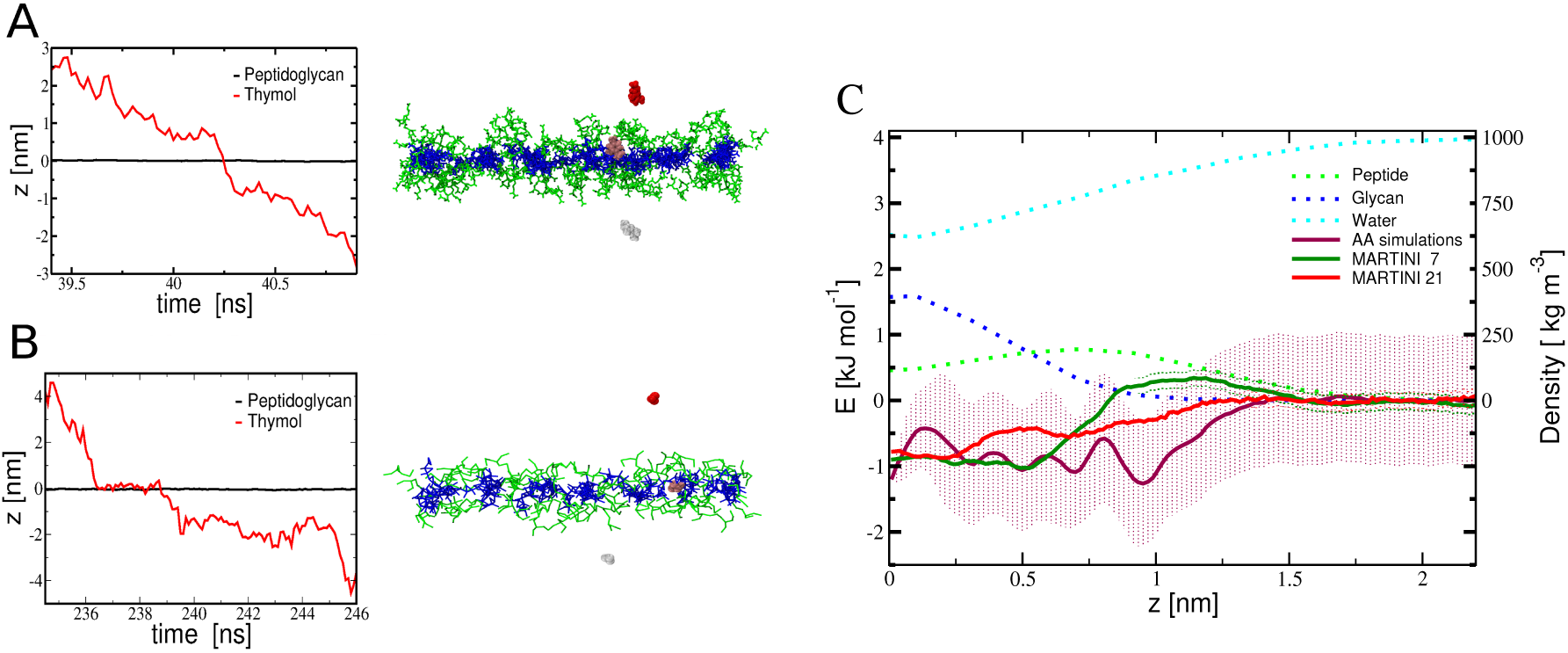
Simulation trajectories and snapshots showing a translocation event in restraint-free AA (A) and MARTINI (B) simulations. The center of mass of peptidoglycan (black) is located on a plane at z=0, and the center of mass of thymol (red) evolves during the translocation of the molecule. The thymol molecule in the snapshots depict three instances - before translocation (red), in the membrane (pink) and after translocation (white). Water is not shown for clarity. (C) The potential of mean force for thymol interacting with peptidoglycan networks along the reaction co-ordinate *z*. The MARTINI profiles for PG network-1 (green) and PG network-2 (red) seem to accurately capture the free energy difference obtained in the all-atom simulations (maroon). The dotted lines indicate density profiles for sugar strands (blue), peptides (green) and water (cyan).

Since the glycan strands for the PG network-1 are tethered with weak restraints in umbrella sampling simulations due to the relatively smaller patch size with no linkages across the periodic boundaries, the PMF calculations were repeated in MARTINI simulations using the larger system comprised of 21 glycan strands (PG network-2), wherein the glycan strands as well as the peptides are covalently bonded with the nearest periodic images. The PG network-2 in AA description, which contains nearly half a million interacting sites, is computationally intractable for PMF calculations by umbrella samplings and hence we did not pursue this here.

As evident in Figure 12C, the free energy profile in MARTINI simulations for PG network-1 matches fairly well with its all-atom profile, except for a small difference of ∼ 1.5 kJ/mol at ∼ 1 nm located in the vicinity of the peptide residues. The appearance of a weak minima in all-atom profile at the locations of peptides is absent in MARTINI simulations. The free energy profile for the MARTINI PG network-2 showed no significant differences from that of the smaller size PG network-1. To sum up, the observation of frequent translocation of thymol in the restraint-free simulations, together with the free energy profiles, support the fact that thymol finds no significant barrier while traversing through peptidoglycan.

## 4 Conclusions

The present work proposes a MARTINI model of bacterial cell wall. The model is parametrized by mapping the distributions for bonded interactions with the virtual CG distributions obtained from all-atom simulations. The aggregation issue for MARTINI sugars is resolved by scaling the energy parameter for non-bonded interactions. As summarized in Table 8, the model reproduces structural properties such as end-to-end distance for glycan strand, the native periodicity of peptide orientation, and area per disaccharides. The mechanical properties from the MARTINI description were compared with that obtained from all-atom data. The response to areal expansion against lateral stresses is in good agreement with values reported in literature. The bending modulus for the model peptidoglycan is ∼ 1 k_B_T. Insertion free energies indicate that the peptidoglycan network offers no resistance for thymol permeation.

In summary we have successfully developed a MARTINI model for peptidoglycan which can be used to reliably assess the structural, mechanical properties as well as the insertion free energies for small molecules. It would be interesting to develop a multilayered model for peptidoglycan using the MARTINI parameters developed here and assess the changes to mechanical properties as well as insertion free energies. We expect our model to have a direct bearing on understanding the barrier properties for the peptidoglycan in Gram-negative bacteria. Furthermore, the proposed CG model will be useful in simulating phenomena associated with bacterial cell walls at larger length and time scales, overcoming the limitations of AA models.

## ASSOCIATED CONTENT

### Supporting Information

List of itp files including force field for non-bonded interactions, a simulation movie for thymol permeation, CG mapping scheme for thymol, histograms for umbrella sampling biasing potential, simulation snapshots for glycan strand at varying scaling factor, parametric optimization algorithm, Figures for bonded distributions for 16-mer glycan strands, and simulation snapshots for peptidoglycan network under stressed conditions.

## Acknowledgement

We acknowledge Unilever Research and Development (Bengaluru, India) for a research grant. We also thank the Supercomputer Education and Research Center (SERC) and Thematic Unit of Excellence on Computational Materials Science (TUE-CMS) - a Department of Science and Technology (DST) supported facility at the Indian Institute of Science Bangalore. We also thank Durba Sengupta for initial discussions on the MARTINI model.

---

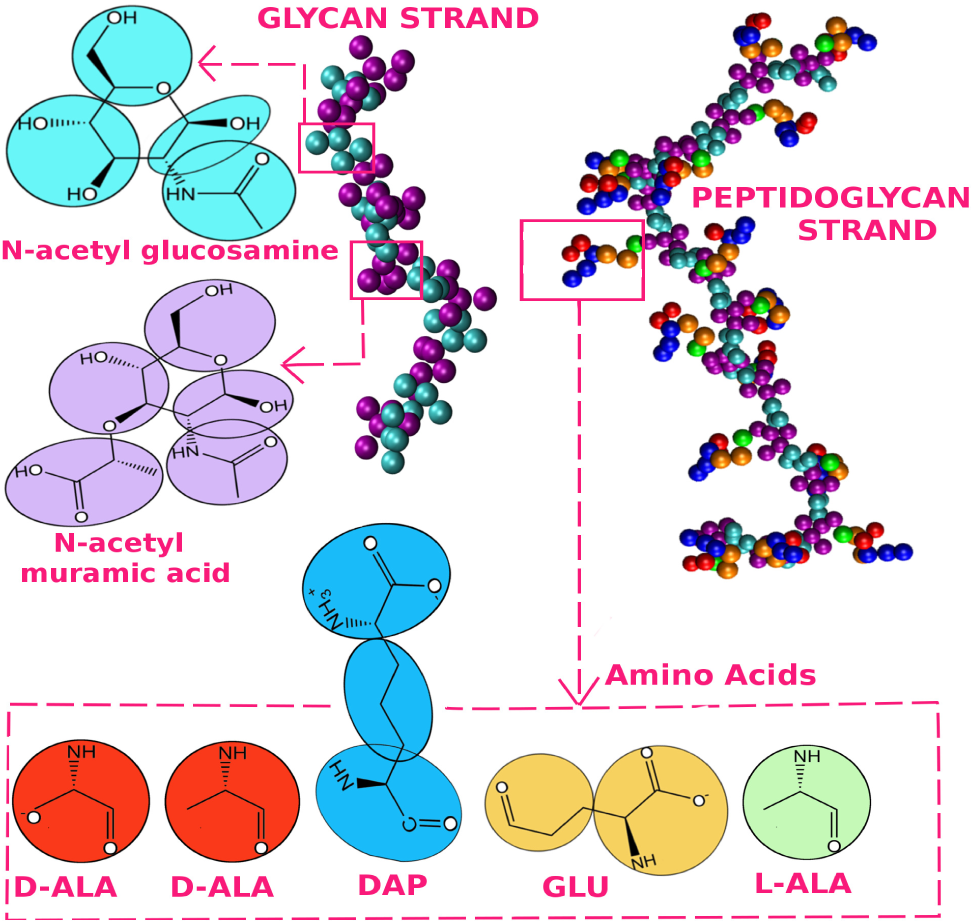
Table of Contents (TOC) graphic

