## Supplementary material for "Developing a Coarse-Grained Model for Bacterial Cell Walls and Evaluating Mechanical Properties and Free Energy Barriers": SI

Supporting Information

Rakesh Vaiwala,[a] Pradyumn Sharma,[a] Mrinalini Puranik [b], K. Ganapathy Ayappa [a,c]

[a] *Department of Chemical Engineering, Indian Institute of Science, 560012 Bangalore, India*
[b] *Unilever Research & Development, 64 Main Road, Whitefield, Bangalore 560066, India*
[c] *Centre for BioSystems Science and Engineering, Indian Institute of Science, 560012 Bangalore, India*

1. All-atom molecular topology for glycan strand comprised of 8 disaccharides:
   AA-glycanstrand-8mer.itp
2. All-atom molecular topology for peptidoglycan strand comprised of 8 disaccharides:
   AA-peptidoglycanstrand-8mer.itp
3. MARTINI molecular topology for glycan strand comprised of 8 disaccharides:
   MARTINI-glycanstrand-8mer.itp
4. MARTINI molecular topology for peptidoglycan strand comprised of 8 disaccharides:
   MARTINI-peptidoglycanstrand-8mer.itp
5. MARTINI force field file for non-bonded interactions:
   martini-forcefield.itp
6. A simulation movie showing permeation of thymol with peptidoglycan network comprised of 7 glycan strands (PG network-1) in a restraint-free MARTINI simulation:
   simulation-movie-peptidoglycan-thymol-7strands.mpg

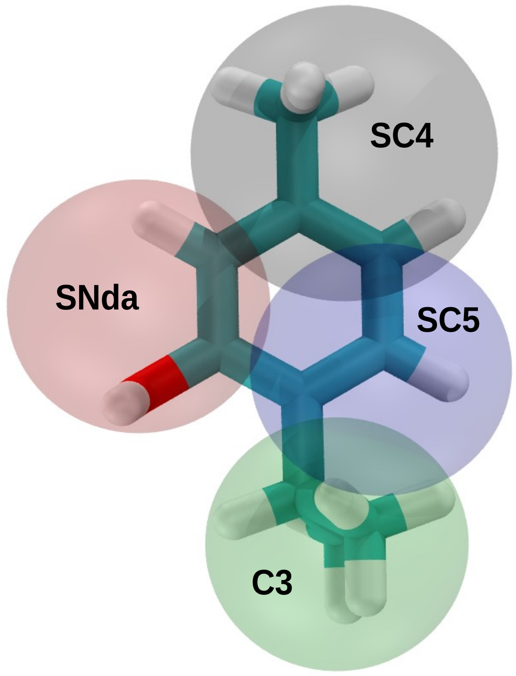


**Figure S1** Licorice representation of all-atom model of thymol, with superimposed translucent coarse-grained beads showing the mapping scheme. The labels indicate the MARTINI bead types.

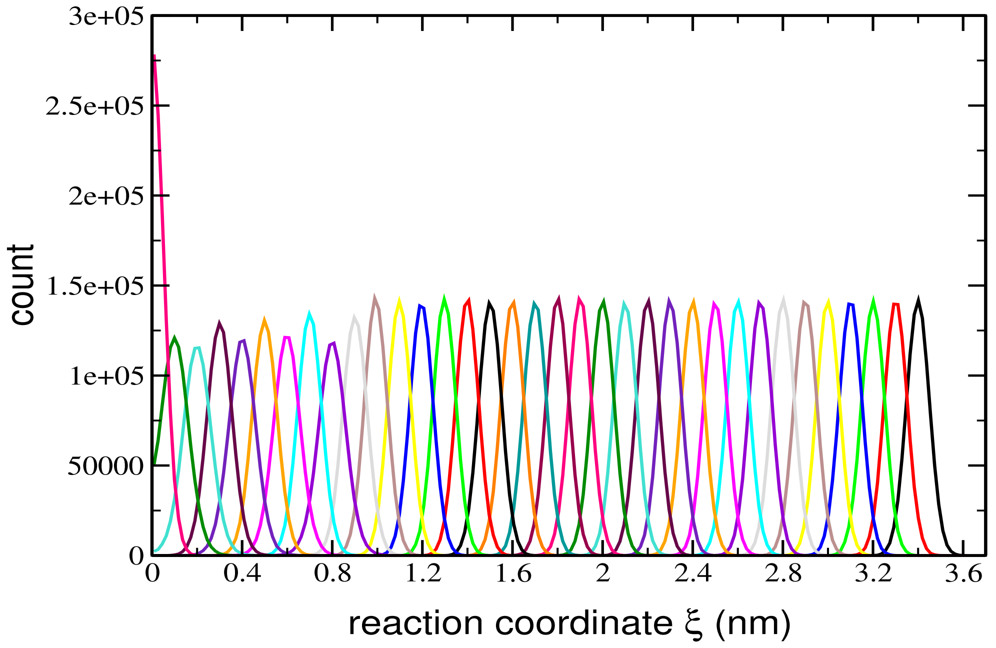


**Figure S2** Histograms for umbrella sampling potential across the reaction coordinate, which is the distance between center of mass of thymol and that of peptidoglycan sheet comprising 7 glycan strands in AA simulations.

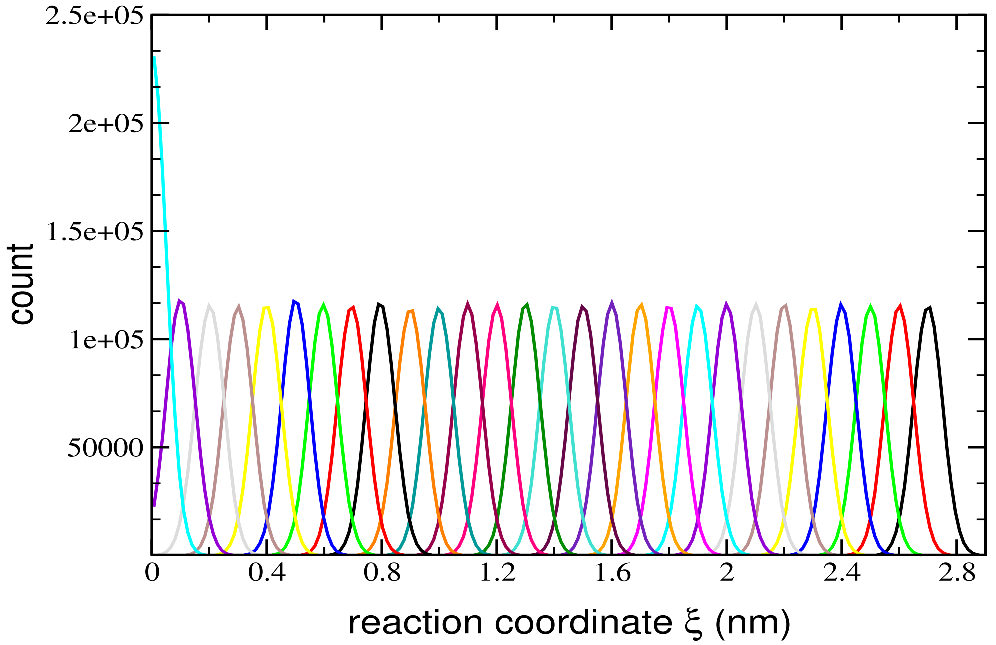


**Figure S3** Histograms for umbrella sampling potential across the reaction coordinate, which is the distance between center of mass of thymol and that of peptidoglycan sheet comprising 7 glycan strands in MARTINI simulations.

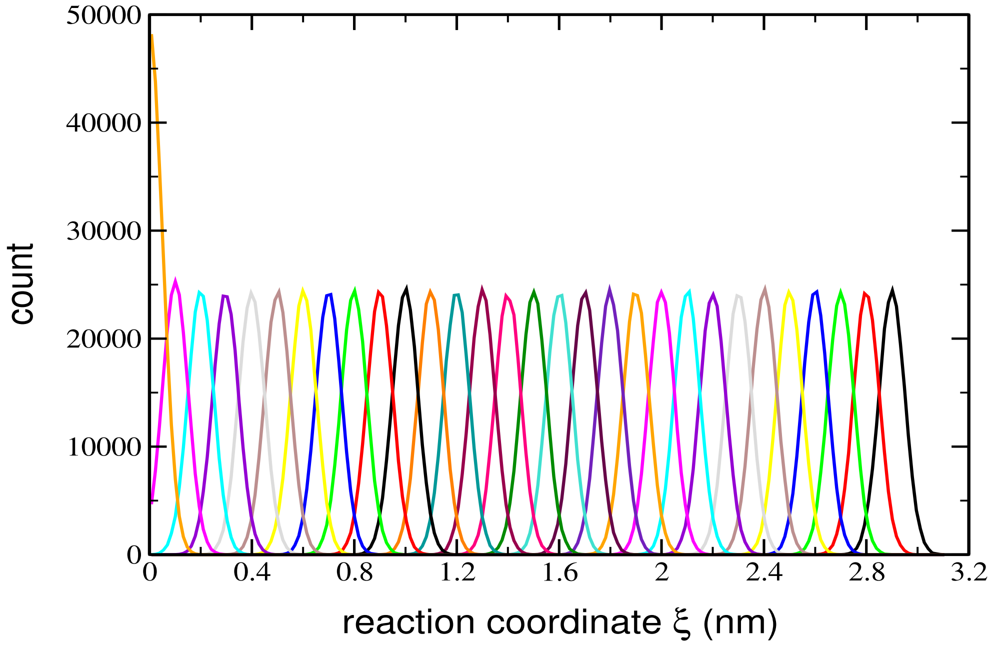


**Figure S4** Histograms for umbrella sampling potential across the reaction coordinate, which is the distance between center of mass of thymol and that of peptidoglycan sheet comprising 21 glycan strands in MARTINI simulations.

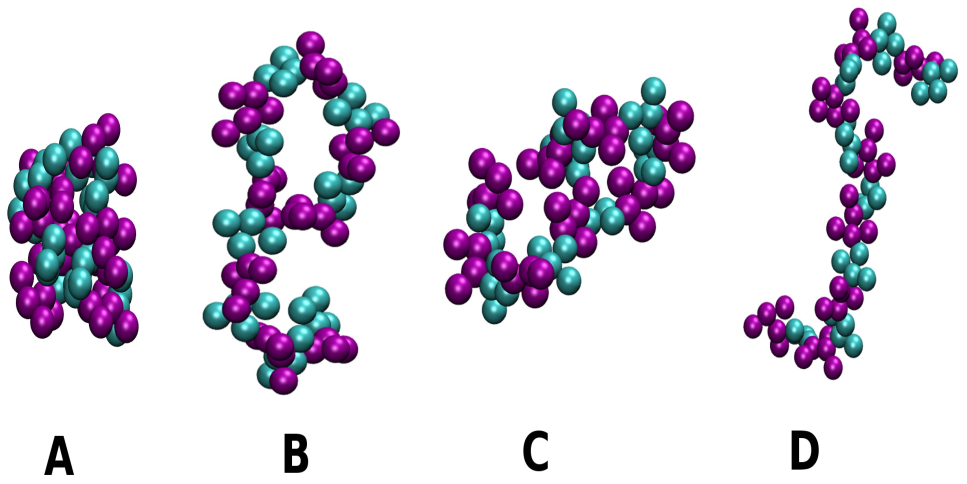


**Figure S5** Glycan strand with scaling factor ($\alpha$) indicating (A) excessive bead aggregation without scaling, (B) with $\alpha = 0.9$, (C) $\alpha = 0.8$ and (D) optimized $\alpha = 0.7$.

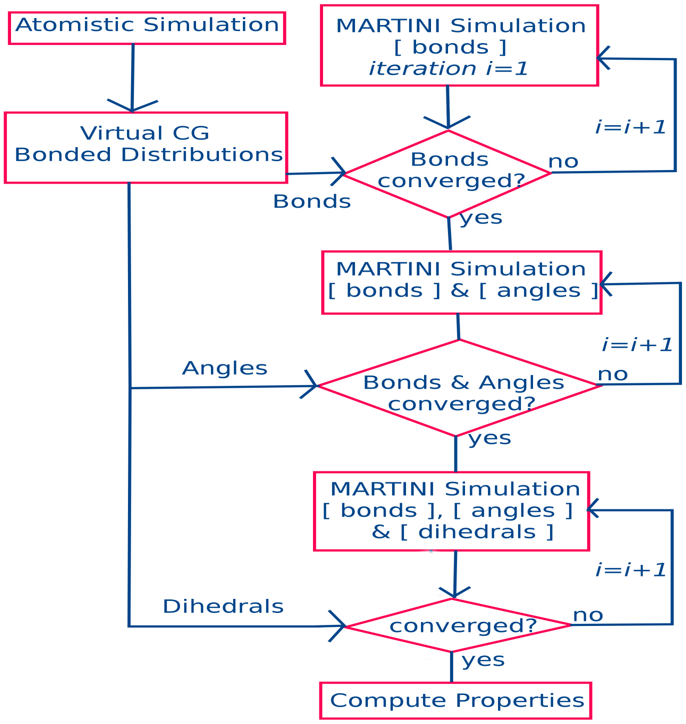


**Figure S6** Algorithm depicting parametric optimization procedure for the bonded interactions.

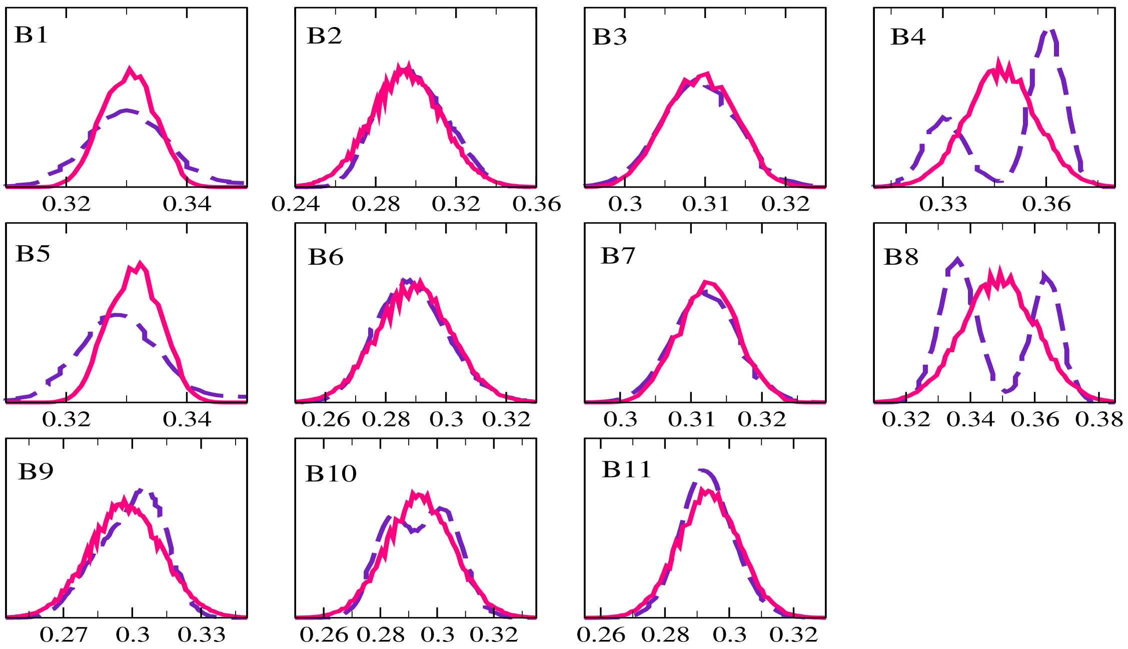


**Figure S7** Mapping of bond distributions in MARTINI simulations with AA target distributions for a 16-mer glycan strand. Solid lines correspond to distributions in MARTINI simulations, and dotted lines indicate distributions in AA simulations. The bond lengths are in nm. The bond labels are according to Table 2 in the article.

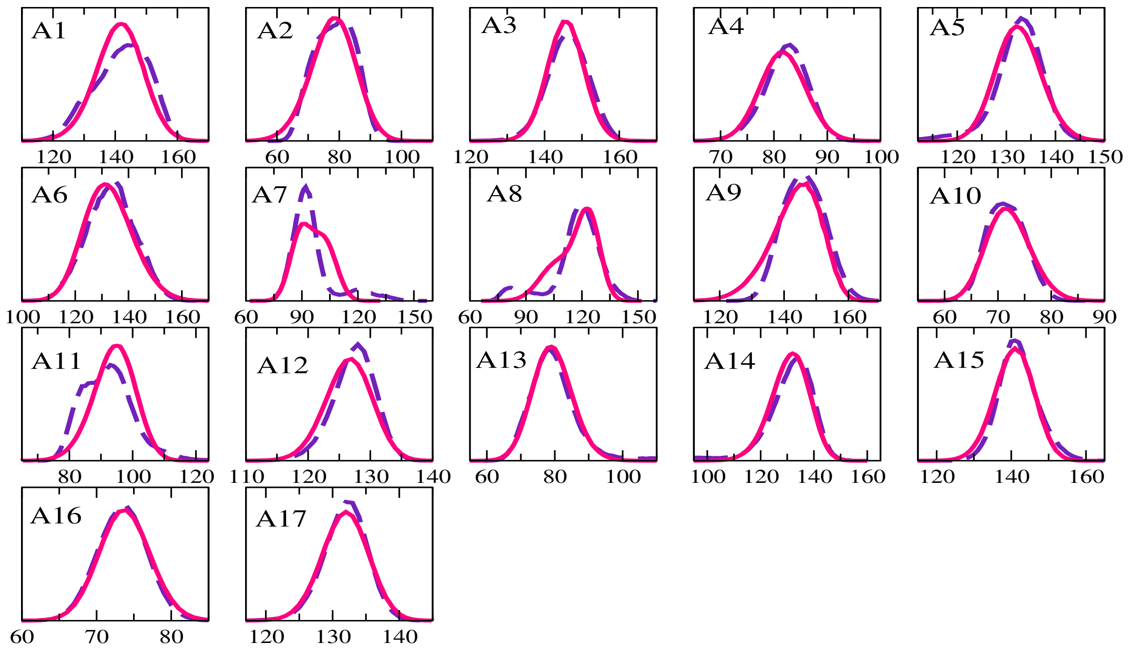


**Figure S8** Mapping of angle distributions in MARTINI simulations with AA target distributions for a 16-mer glycan strand. Solid lines correspond to distributions in MARTINI simulations, and dotted lines indicate distributions in AA simulations. The angles are in degree. The lables for angles are according to Table 3 in the article.

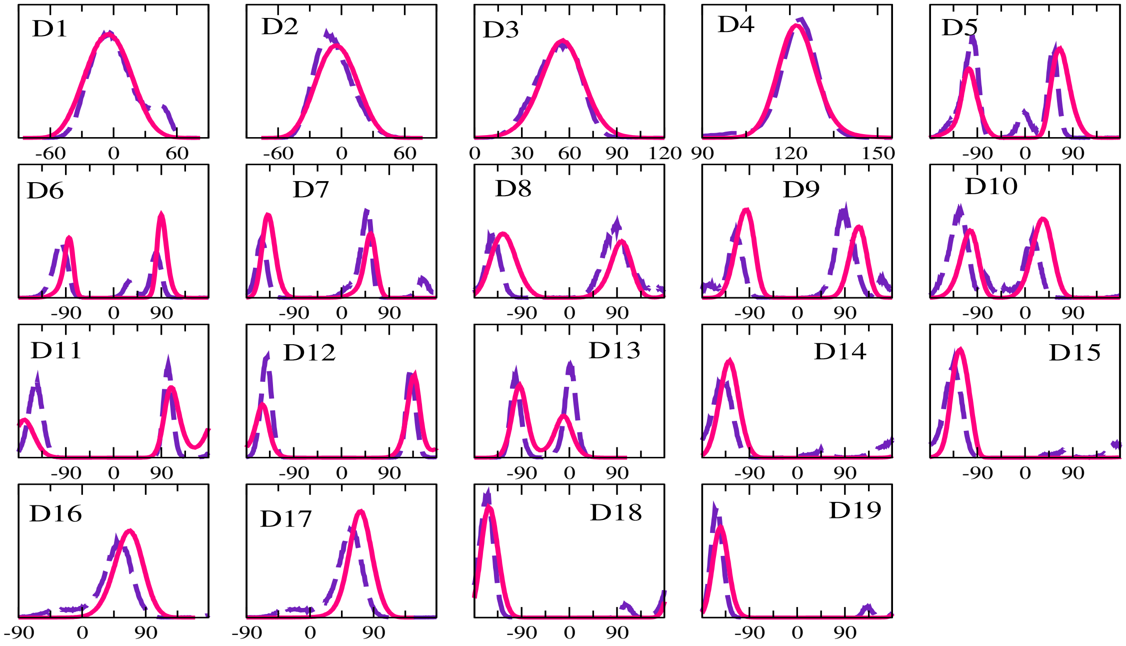


**Figure S9** Mapping of dihedral angle distributions in MARTINI simulations with AA target distributions for a 16-mer glycan strand. Solid lines correspond to distributions in MARTINI simulations, and dotted lines indicate distributions in AA simulations. The angles are in degree. The labels for dihedrals are according to Table 4 in the article.

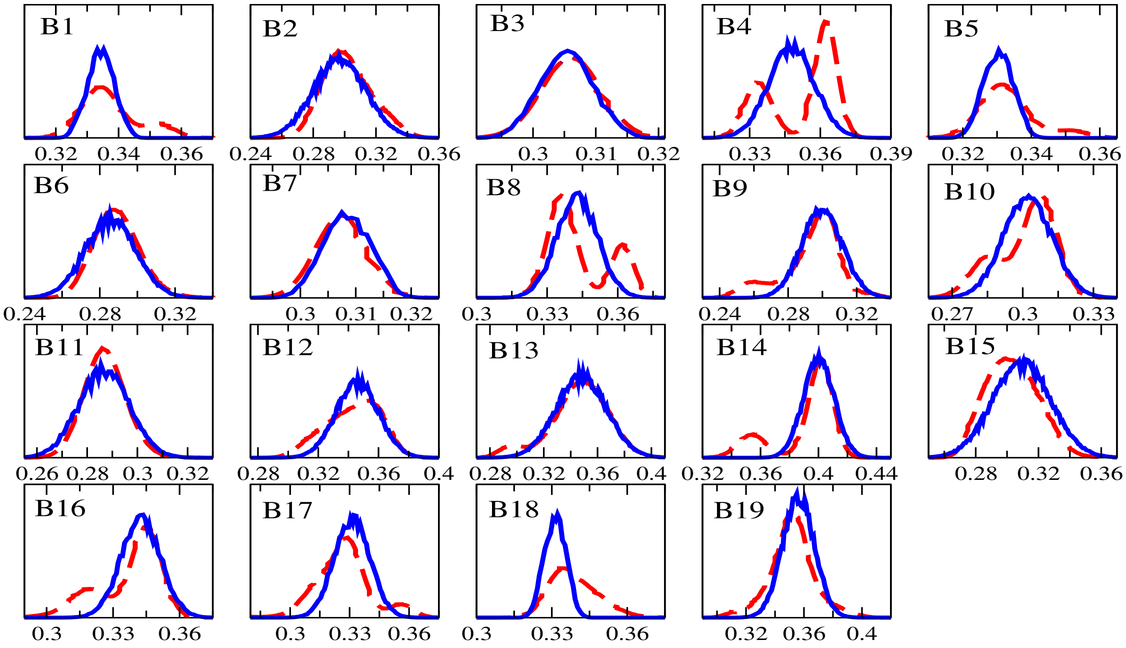


**Figure S10** Mapping of bond distributions in MARTINI simulations with AA target distributions for a 16-mer peptidoglycan strand. Solid lines correspond to distributions in MARTINI simulations, and dotted lines indicate distributions in AA simulations. The bond lengths are in nm. The bond labels are according to Table 5 in the article.

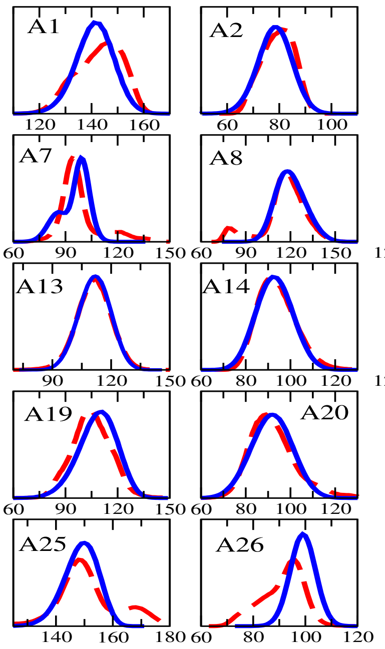


**Figure S11** Mapping of angle distributions in MARTINI simulations with AA target distributions for a 16-mer peptidoglycan strand. Solid lines correspond to distributions in MARTINI simulations, and dotted lines indicate distributions in AA simulations. The angles are in degree. The labels for angles are according to Table 6 in the article.

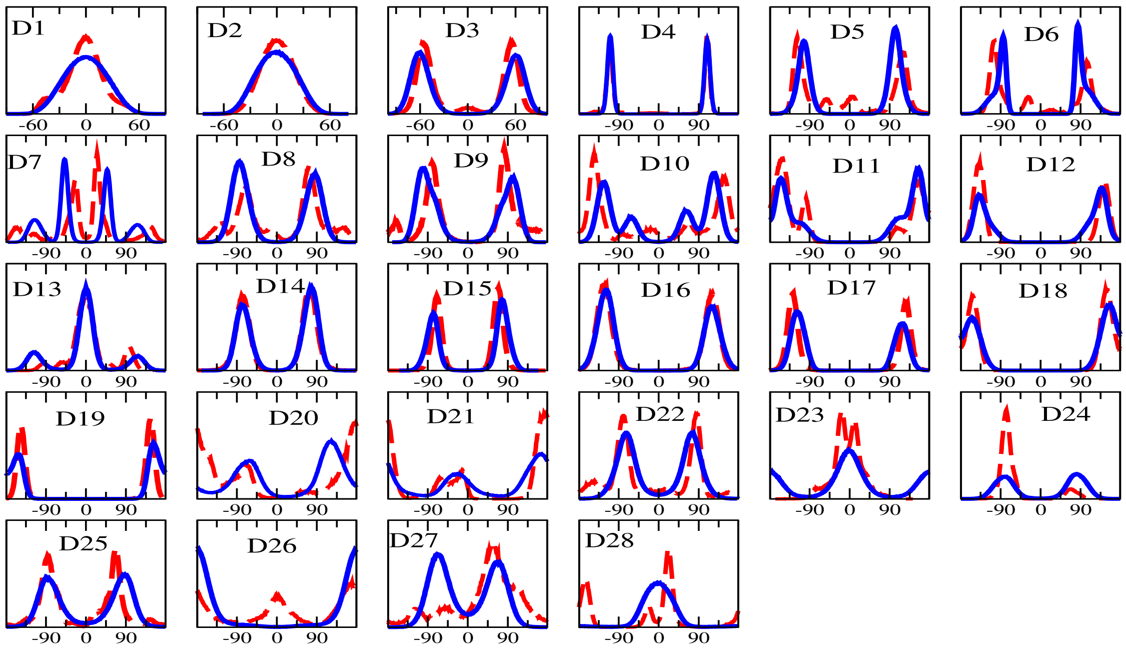


**Figure S12** Mapping of dihedral angle distributions in MARTINI simulations with AA target distributions for a 16-mer peptidoglycan strand. Solid lines correspond to distributions in MARTINI simulations, and dotted lines indicate distributions in AA simulations. The angles are in degree. The labels for diherdals are according to Table 7 in the article.

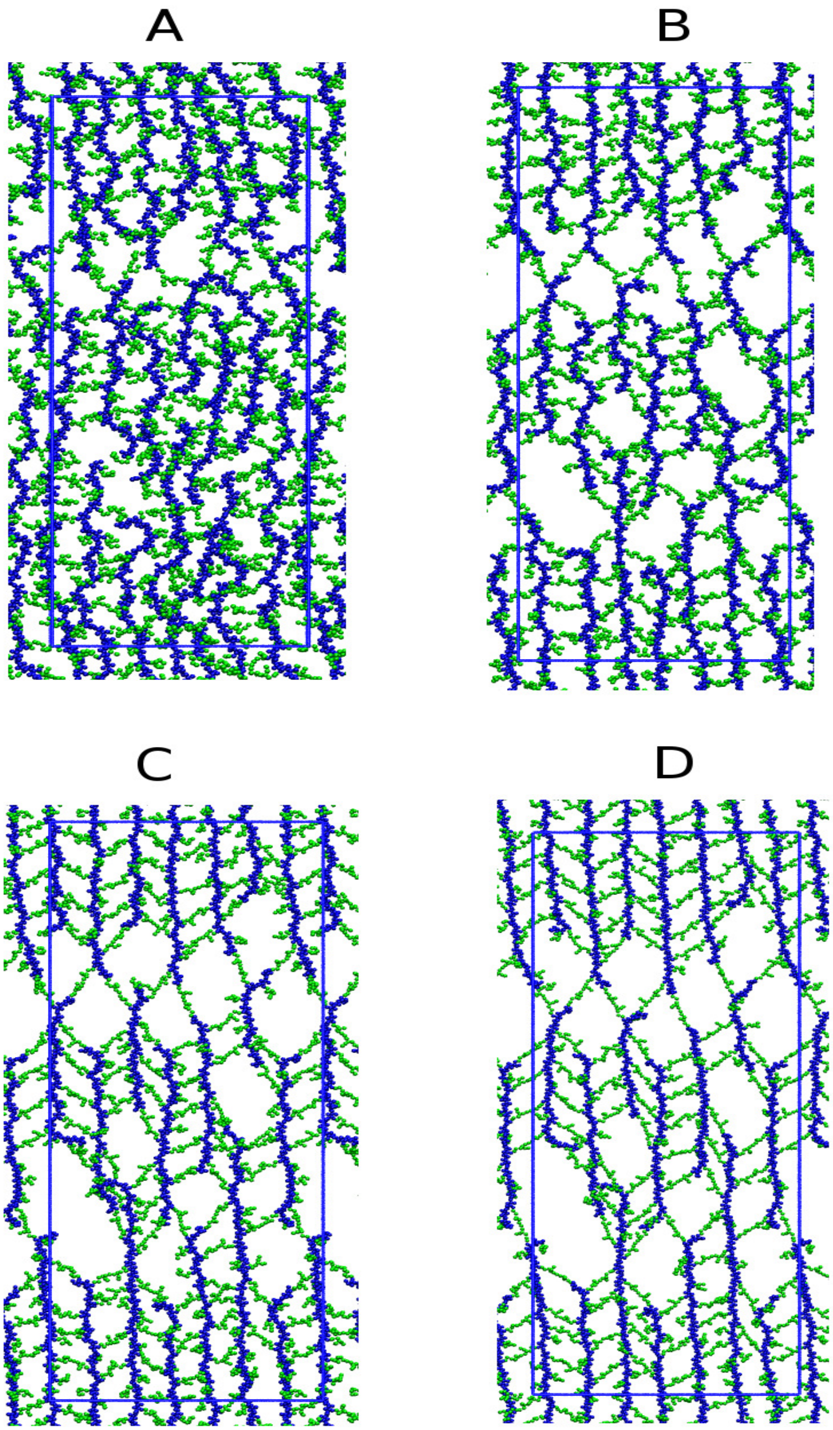


**Figure S13** Simulation snapshots showing the peptidoglycan PG network-2 (21 glycan strands) in its (A) relax state and (B-D) stressed conditions. The tension values and corresponding areal expansion are (B) 8.18 mN/m and 60.5%, (C) 20.09 mN/m and 87.6%, and (D) 49.27 mN/m and 117%. The glycan strands are indicated by blue color, and the peptides are shown in green. Water and ions are not shown for visual clarity. The rectangular box shows the central simulation cell. The peptide linkages and the continuity of glycan strands across the periodic boundaries are also depicted.
